# SARS-CoV-2 Spike Glycoprotein and ACE2 interaction reveals modulation of viral entry in wild and domestic animals

**DOI:** 10.1101/2020.05.08.084327

**Authors:** Manas Ranjan Praharaj, Priyanka Garg, Veerbhan Kesarwani, Neelam A Topno, Raja Ishaq Nabi Khan, Shailesh Sharma, Manjit Panigrahi, B P Mishra, Bina Mishra, G Sai kumar, Ravi Kumar Gandham, Raj Kumar Singh, Subeer Majumdar, Trilochan Mohapatra

## Abstract

**Background:** SARS-CoV-2 is a viral pathogen causing life-threatening disease in human. Interaction between spike protein of SARS-CoV-2 and ACE2 receptor on the cells is a potential factor in the infectivity of a host. The interaction of SARS-CoV-2 spike receptor-binding domain with its receptor - ACE2, in different hosts was evaluated to understand and predict viral entry. The protein and nucleotide sequences of ACE2 were initially compared across different species to identify key differences among them. The ACE2 receptor of various species was homology modeled (6LZG, 6M0J, and 6VW1 as a reference), and its binding ability to the spike ACE2 binding domain of SARS-CoV-2 was assessed. Initially, the spike binding parameters of ACE2 of known infected and uninfected species were compared with each Order (of animals) as a group. Finally, a logistic regression model vis-a-vis the spike binding parameters of ACE2 (considering data against 6LZG and 6M0J) was constructed to predict the probability of viral entry in different hosts.

**Results:** Phylogeny and alignment comparison did not lead to any meaningful conclusion on viral entry in different hosts. Out of several spike binding parameters of ACE2, a significant difference between the known infected and uninfected species was observed for six parameters. However, these parameters did not specifically categorize the Orders (of animals) into infected or uninfected. The logistic regression model constructed revealed that in the mammalian class, most of the species of Carnivores, Artiodactyls, Perissodactyls, Pholidota, and Primates had high probability of viral entry. However, among the primates, African Elephant had low probability of viral entry. Among rodents, hamsters were highly probable for viral entry with rats and mice having a medium to low probability. Rabbits have a high probability of viral entry. In Birds, ducks have a very low probability, while chickens seemed to have medium probability and turkey showed the highest probability of viral entry.

**Conclusions:** Most of the species considered in this study showed high probability of viral entry. This study would prompt us to closely follow certain species of animals for determining pathogenic insult by SARS-CoV-2 and for determining their ability to act as a carrier and/or disseminator.

## Background

Three large-scale disease outbreaks during the past two decades, *viz.,* Severe Acute Respiratory Syndrome (SARS), Middle East Respiratory Syndrome (MERS), and Swine Acute Diarrhea Syndrome (SADS) were caused by three zoonotic coronaviruses. SARS and MERS, which emerged in 2003 and 2012, respectively, caused a worldwide pandemic claiming 774 (8,000 SARS cases) and 866 (2,519 MERS cases) human lives, respectively[1], while SADS devastated livestock production by causing fatal disease in pigs in 2017. The SARS and MERS viruses had several common factors in having originated from bats in China and being pathogenic to human or livestock[2–4]. Seventeen years after the first highly pathogenic human coronavirus, SARS-COV-2 is devastating the world with 87,808,867 cases and 1,894,632 deaths (as on Jan 07, 2021)[5]. This outbreak was first identified in Wuhan City, Hubei Province, China, in December 2019 and notified by WHO on 5^th^ January 2020. The disease has since been named as COVID-19 by WHO.

Coronaviruses (CoVs) are an enveloped, crown-like viral particles belonging to the subfamily Orthocoronavirinae in the family Coronaviridae and the Order Nidovirales. They harbor a positive-sense, single-strand RNA (+ssRNA) genome of 27–32 kb in size. Two large overlapping polyproteins, ORF1a and ORF1b, that are processed into the viral polymerase (RdRp) and other nonstructural proteins involved in RNA synthesis or host response modulation, cover two thirds of the genome. The rest 1/3 of the genome encodes for four structural proteins (spike (S), envelope (E), membrane (M), and nucleocapsid (N)) and other accessory proteins. The four structural proteins and the ORF1a/ORF1b are relatively consistent among the CoVs, however, number and size of accessory proteins govern the length of the CoV genome[4]. This genome expansion is said to have facilitated acquisition of genes that encode accessory proteins, which are beneficial for CoVs to adapt to a specific host[6, 7]. Next generation sequencing has increased the detection and identification of new CoV species resulting in expansion of CoV subfamily. Currently, there are four genera (α-, β-, δ-, and γ-) with thirty-eight unique species in CoV subfamily (ICTV classification) including the three highly pathogenic CoVs, *viz.,* SARS-CoV-1, MERS-CoV, SARS-CoV-2 are β-CoVs[8].

Coronaviruses are notoriously promiscuous. Bats host thousands of these types, without succumbing to illness. The CoVs are known to infect mammals and birds, including dogs, chickens, cattle, pigs, cats, pangolins, and bats. These viruses have the potential to leap to new species and in this process mutate along the way to adapt to their new host(s). COVID-19, global crisis likely started with CoV infected horseshoe bat in China. The SARS-CoV-2 is spreading around the world in the hunt of entirely new reservoir hosts for re-infecting people in the future[9]. Recent reports of COVID-19 in a Pomeranian dog and a German shepherd in Hong Kong[10]; in a domestic cat in Belgium[11]; in five Malayan tigers and three lions at the Bronx Zoo in New York City[12] and in minks[13] make it all the more necessary to predict species that could be the most likely potential reservoir hosts in times to come.

Angiotensin-converting enzyme 2 (ACE2), an enzyme that physiologically counters RAAS activation functions as a receptor for both the SARS viruses (SARS-CoV-1 and SARS-CoV-2)[14–16]. The ACE2 human RefSeqGene is 48037 bp in length with18 exons and is located on chromosome X. ACE2 is found attached to the outer surface of cells in the lungs, arteries, heart, kidney, and intestines[17, 18]. The potential factor in the infectivity of a cell is the interaction between SARS viruses and the ACE2 receptor[19, 20]. By comparing the ACE2 sequence, several species that might be infected with SARS-CoV2 have been identified[21]. R
ecent studies, exposing cells/animals to the SARS-CoV2, revealed humans, horseshoe bats, civets, ferrets, cats and pigs could be infected with the virus and mice, dogs, pigs, chickens, and ducks could not be or poorly infected[16, 22]. Pigs, chickens, fruit bats, and ferrets are being exposed to SARS-CoV2 at Friedrich-Loeffler Institute and initial results suggest that Egyptian fruit bats and ferrets are susceptible, whereas pigs and chickens are not[23]. In this cause of predicting potential hosts, no studies on ACE2 sequence comparison among species along with homology modeling and prediction, to define its interaction with the spike protein of SARS-CoV-2 are available. Therefore, the present study is taken to identify viral entry in potential hosts through sequence comparison, homology modeling and prediction.

## Results

### Sequence comparison of ACE2

The protein and DNA sequence lengths of ACE2 varied in different hosts (Supplementary Table 1). Among the sequences that were compared, the longest CDS was found in the Order - Chiroptera *(Myotis brandtii* - 811 aa) and the smallest in the Order – Proboscidea (*Loxodonta africana* - 800 aa). The within group mean distance, the parameter indicative of variability of nucleotide sequences within the group was found to be minimum in Perrisodactyla followed by Primates and was maximum among the Galliformes followed by Chiroptera (Supplementary Table 2). To establish the probability of SARS-CoV-2 entry into species of other Orders, the distance of all Orders from Primates was assessed (Supplementary Table 3). This distance was found minimum for Perissodactyls followed by Carnivores and maximum for Galliformes followed by Anseriformes. Further, to decide a cut-off distance that can establish whether the species can be infected or not, the individual distance of each species from *Homo sapiens* was evaluated (Supplementary Table 3). *Meleagris gallopavo* (Turkey) is the species, which had the greatest distance from *Homo sapiens*. The minimum distance that corresponded to the species that was already established to be uninfected with the SARS-CoV-2 i.e. *Sus scrofa,* was 0.194. The codon-based test of neutrality to understand the selection pressure on the ACE2 sequence in the process of evolution was done. The analysis showed that there was a significant negative selection between and within Orders for the ACE2 sequence. On sequence comparison of the spike interacting domain the alignments, both protein and nucleotide (Supplementary Figure 1 and 2) showed that the sequences were well conserved within the Orders, suggesting that the structure defined by the sequence was conserved within the Orders. The maximum variability with the *Homo sapiens* sequence within these regions was observed for Galliformes, followed by Accipitriformes, Testudines, Crocodilia and Chiroptera. The protein sequence alignment at 30-41aa, 82-84 aa and 353-357 also showed similar sequence conservation and variability.

### Phylogenetic analysis

The protein sequences aligned were further subjected to find the best substitution model for phylogenetic analysis. The best model on the basis of BIC was found to be JTT + G. The phylogenetic analysis clearly classified the sequences of the species into their Orders. All the sequences were clearly grouped into two clusters. The first cluster represented the Mammalian class and the second cluster was represented by two sub-clusters of Avian and Reptilian classes with high bootstrap values (Figure 1). Within the mammalian cluster, the artiodactyls were sub-clustered farthest to the primates and the rodents, lagomorphs and carnivores were found clustered close to the primates with reliable bootstrap values. The Chiroptera sub-cluster had a sub-node constituting horseshoe bat *(Rhinolophus ferrumequinum)* and the fruit bats *(Pteropus Alecto and Rousettus aegyptiacus)* (Figure 1).

**Figure 1.**
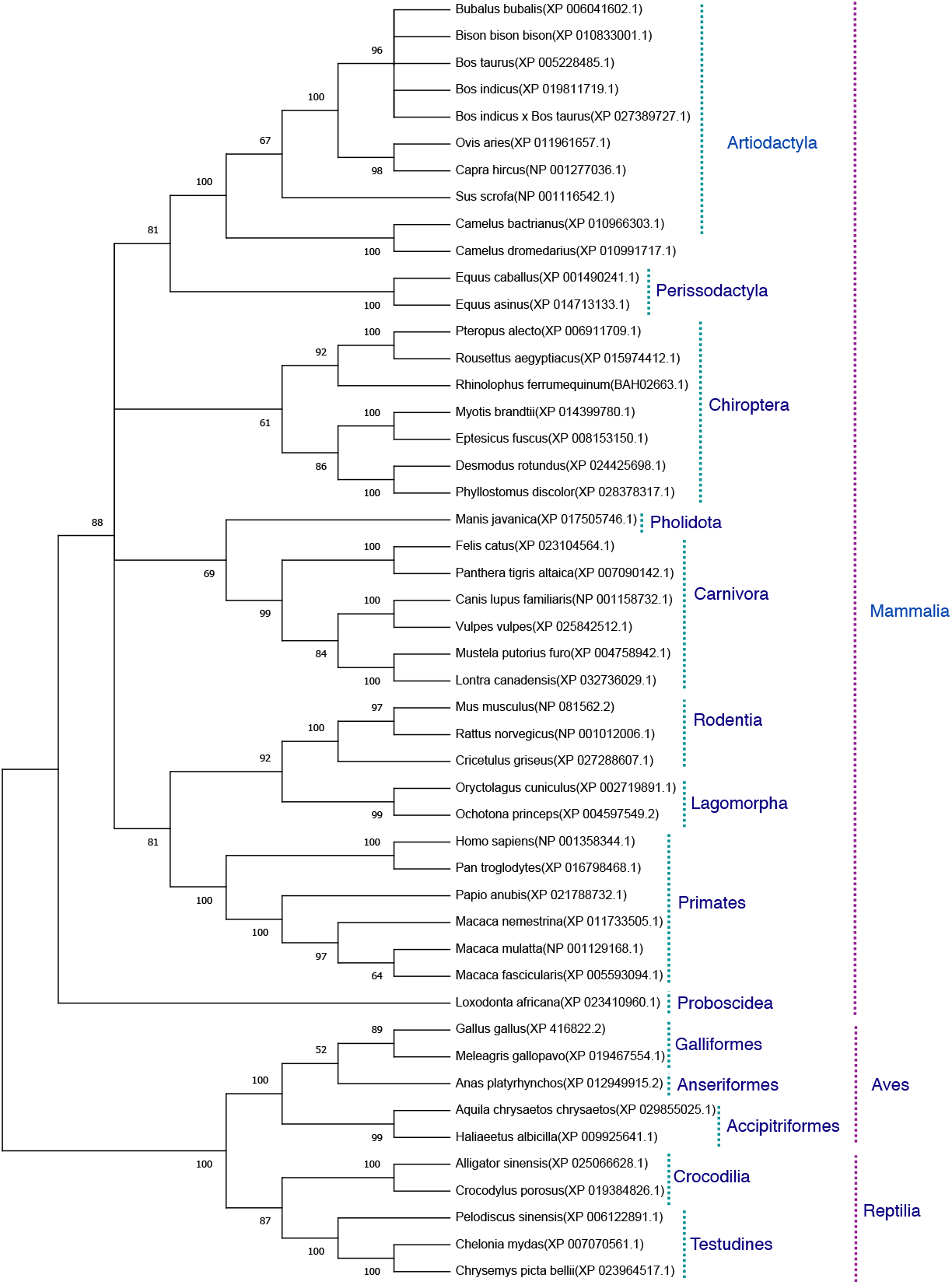
Phylogenetic analysis of ACE2 protein sequences. The tree was constructed using neighbor joining method in MEGA 6.0. The bootstrap values are given at each node.

### Homology modelling, docking and evaluation of spike binding parameters of ACE2

Homology modeling was done for all the ACE2 sequences based on the X-ray diffraction structures defined in PDB database - 6LZG, 6VW1 and 6M0J. After homology modelling using SWISS-MODEL, the models (144 = 48 × 3) were validated using SAVES. The homology modelled structures used in this study showed no “Error” in PROVE. Most of the homology modelled structure had > 90% score in PROCHECK and > 95% score in ERRAT2 showing the models were good enough for further analysis. All the models were assigned “PASS” by Verify 3D (Supplementary Table 4).

These models constructed were then studied for their interaction with the spike ACE2 - binding domains defined in the same IDs using GRAMM-X (Supplementary Table 5). Out of the 5 docked complexes tested for each X-crystallography structure, the best three docked complexes were selected based on the delta G and the number of Hydrogen bonds. Several spike binding parameters for these selected complexes – 432 were generated in FoldX (Supplementary Table 6). Initially, to classify the infected from the uninfected irrespective of the Order(s) unpaired t-test was done. The spike binding parameters – RMSD, delta G, Intraclashes Group1, Van der Waals and Solvation Hydrophobic and entropy sidechain were found to be significantly different in the infected from the uninfected (Supplementary Table 7 & 8). These parameters were further used to classify an Order as infected or uninfected (Supplementary Table 9). None of the parameters could clearly classify the Orders to be infected or uninfected i.e., for RMSD, the Orders - Artiodactyla and Testudines, were significantly different from the infected and uninfected, however, the Order - Chiroptera was significantly different only from the infected (Figure 2, 3 and 4). Similar findings were observed with the rest of significant parameters that were evaluated. This suggested that the use of a single parameter would not help in identifying a species with probable viral entry.

**Figure 2.**
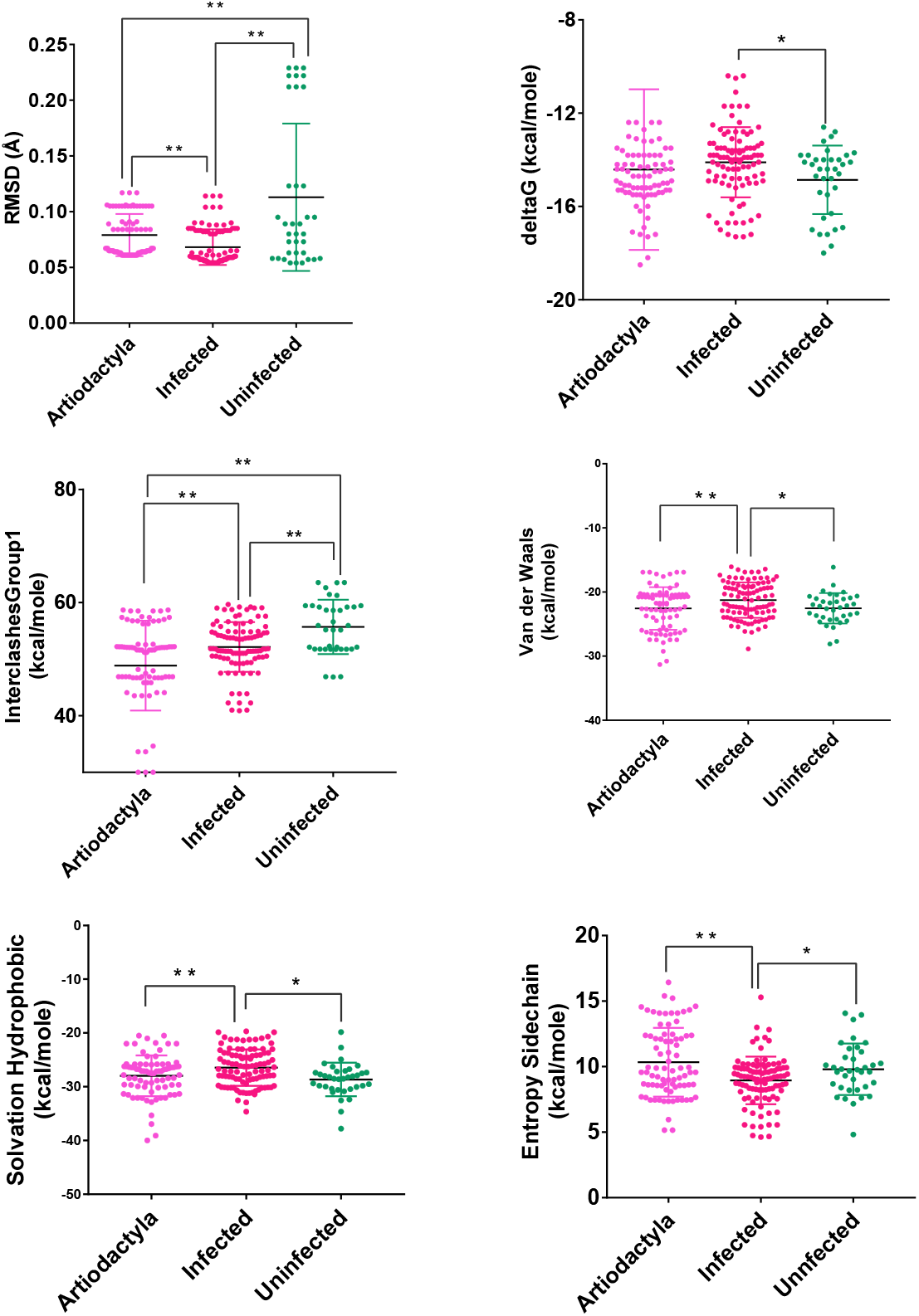
Scatterplot showing the comparison of Artiodactyls with infected and uninfected groups for the all six significant parameters (A). RMSD - Significant difference on comparison of Artiodactyls with infected and uninfected groups. (B). delta G – No significant difference on comparison of Artiodactyls with infected and uninfected groups. (C). InterclashesGroup1 – Significant difference on comparison of Artiodactyls with infected and uninfected groups. (D). Van der Waals – Significant difference on comparison of Artiodactyls with infected and no significant difference with uninfected groups. (E). Solvation hydrophobic - Significant difference on comparison of Artiodactyls with infected and no significant difference with uninfected groups. (F). Entropy side chain - Significant difference on comparison of Artiodactyls with infected group and no significant difference with uninfected group. ** Significance at P < 0.01; * Significance at P < 0.05 after unpaired t test on comparing two groups at a time.

**Figure 3.**
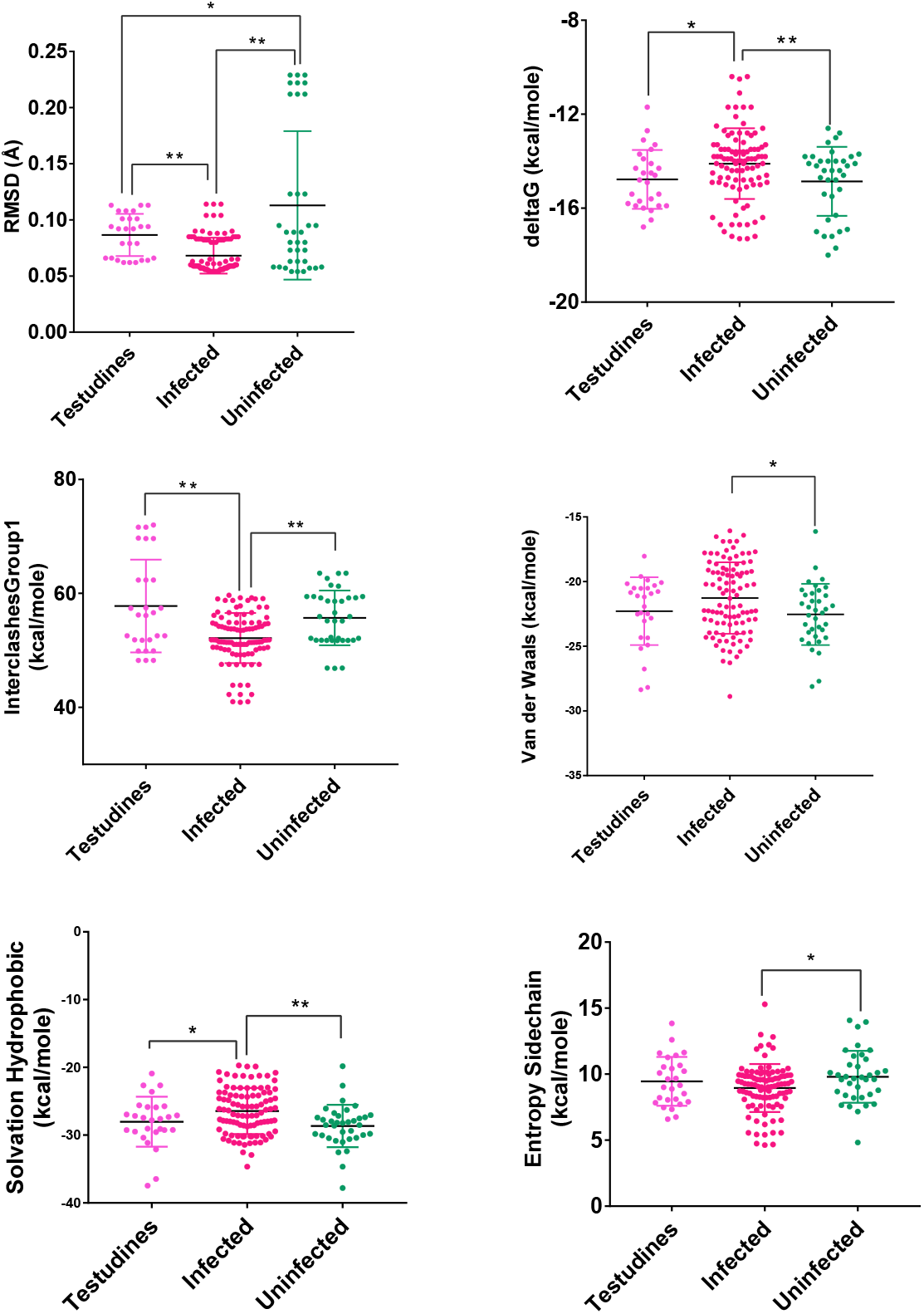
Scatterplot showing the comparison of Testudines with infected and uninfected groups for the all six significant parameters (A). RMSD – Significant difference on comparison of Testudines with infected and uninfected groups. (B). delta G – Significant difference on comparison of Testudines with infected and no significant difference with uninfected groups. (C). InterclashesGroup1 – Significant difference on comparison of Testudines with infected and no significant difference with uninfected groups. (D). Van der Waals – No significant difference on comparison of Testudines with infected and uninfected groups. (E). Solvation hydrophobic - Significant difference on comparison of Testudines with infected and no significant difference with uninfected groups. (F). Entropy side chain – No significant difference on comparison of Testudines with infected group and uninfected group. ** Significance at P < 0.01; * Significance at P < 0.05 after unpaired t test on comparing two groups at a time.

**Figure 4.**
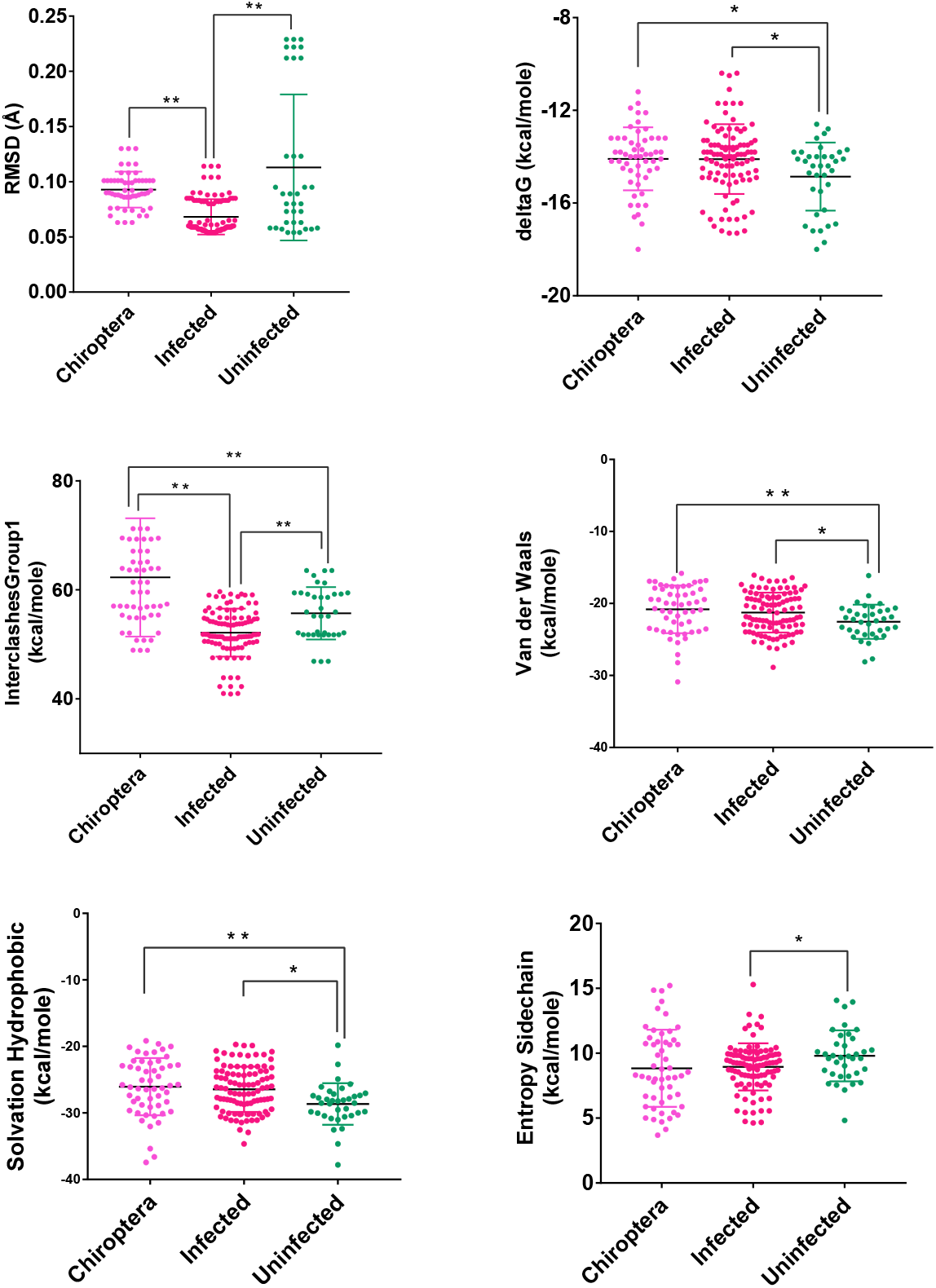
Scatterplot showing the comparison of Chiroptera with infected and uninfected groups for the all six significant parameters (A). RMSD – Significant difference on comparison of Chiroptera with infected and no significant difference with uninfected groups. (B). delta G – Significant difference on comparison of Chiroptera with uninfected and no significant difference with infected groups. (C). InterclashesGroup1 – Significant difference on comparison of Chiroptera with infected and uninfected groups. (D). Van der Waals – Significant difference on comparison of Chiroptera with uninfected and no significant difference with infected groups. (E). Solvation hydrophobic - Significant difference on comparison of Chiroptera with uninfected and no significant difference with infected groups. (F). Entropy side chain – No significant difference on comparison of Chiroptera with infected and uninfected groups. ** Significance at P < 0.01; * Significance at P < 0.05 after unpaired t test on comparing two groups at a time-.

### Logistic regression and prediction of viral entry probability

The seven different combination of data used for finding the best combination of X-Crystallography models for predicting the viral entry can be accessed through supplementary Table 7 (for details please refer to materials and methods). On analyzing the data against a single X-Crystallography model, i.e. either 6M0J or 6LZG or 6VW1, the number of significant parameters at 5% level of significance were found to be highest for 6M0J and lowest for 6VW1 (Table 1). Among these single model combinations, the highest reduction in null deviance and the greatest R square was observed for 6VW1. However, the AIC value was lowest for 6LZG. On considering the data against two models, the number of significant parameters were found to be highest for both the combinations - 6LZG & 6M0J and 6LZG & 6VW1. These two combinations were better than the other combination vis - a - vis most of the evaluation parameters. Between, 6LZG & 6M0J and 6LZG & 6VW1, the former was having the lowest AIC value, the greatest reduction in null deviance and the lowest p-value that determines significant reduction in null deviance than the later. However, the R square was higher in the later than the former. The analysis of data against the three-model combination - 6M0J & 6VW1 & 6LZG, also proved to have good estimates of evaluation parameters (Table 1). Among all the seven data combinations considered, based on the evaluation parameters, the best three combinations - 6LZG & 6M0J and 6LZG & 6VW1 and 6M0J & 6VW1 & 6LZG, were considered for evaluating the probability of viral entry by partitioning the data as training and test data. The predicted probability of all the infected species was closer to being infected with the data combinations - 6M0J & 6LZG followed by 6LZG & 6VW1 and 6M0J & 6VW1 & 6LZG. Similar, was the probability for the uninfected species except for a minor difference in *Sus scrofa.* Considering these findings, the prediction equation obtained from the combination of 6M0J & 6LZG was selected for predicting the probability of the rest of the species in the study. The probabilities were predicted using the following equation: -

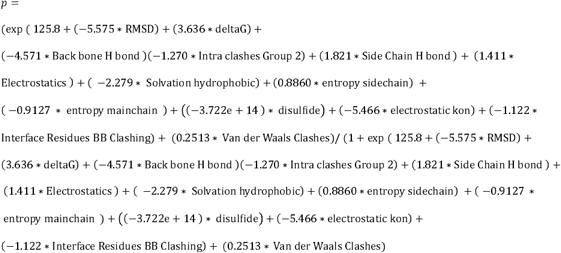

**Table 1:**
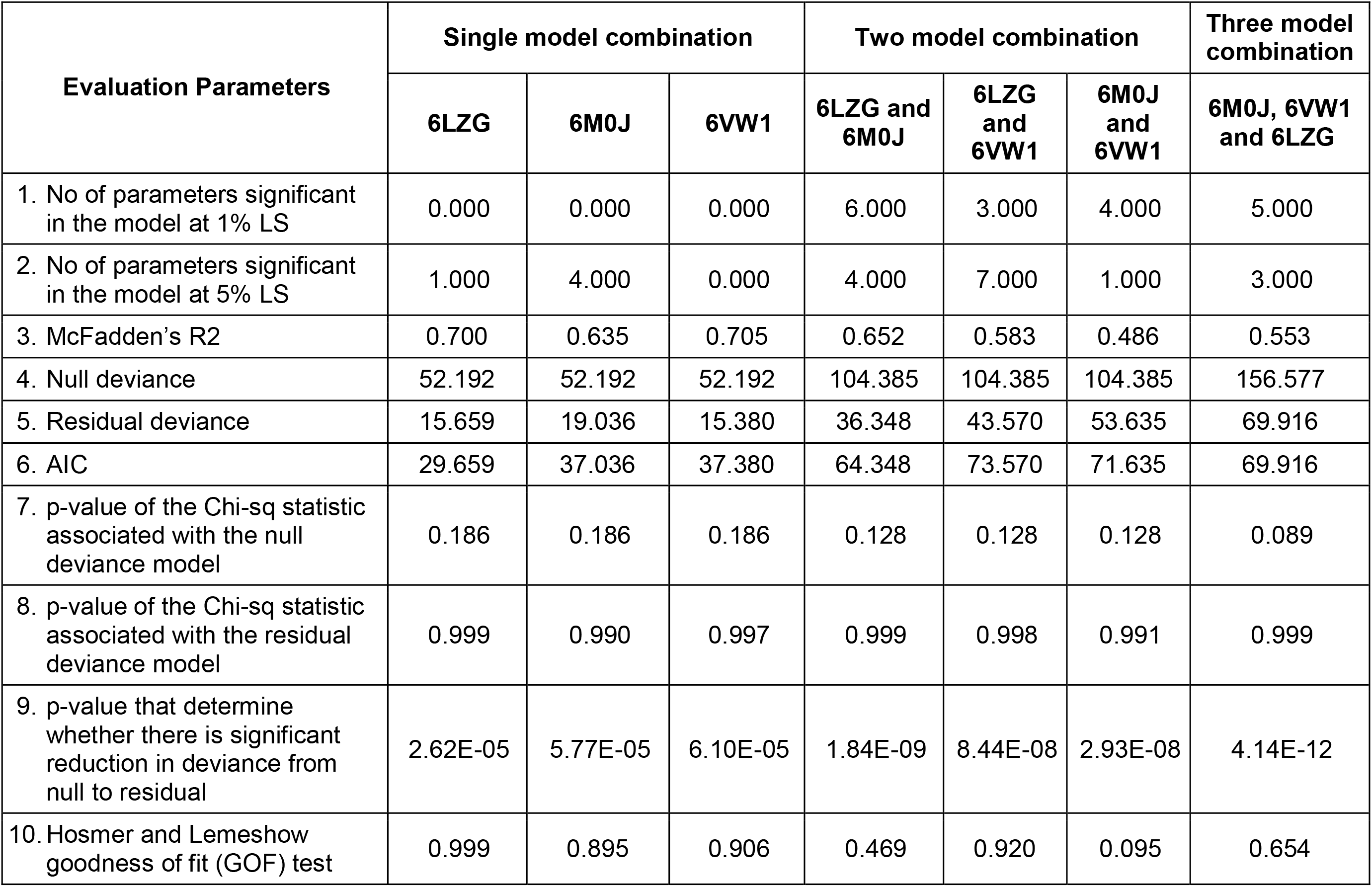
Evaluation of data combinations using logistic regression

| Evaluation Parameters | Single model combination |  |  | Two model combination |  |  | Three model combination |
| --- | --- | --- | --- | --- | --- | --- | --- |
|  | 6LZG | 6M0J | 6VW1 | 6LZG and 6M0J | 6LZG and 6VW1 | 6M0J and 6VW1 | 6M0J, 6VW1 and 6LZG |
| 1. No of parameters significant in the model at 1% LS | 0.000 | 0.000 | 0.000 | 6.000 | 3.000 | 4.000 | 5.000 |
| 2. No of parameters significant in the model at 5% LS | 1.000 | 4.000 | 0.000 | 4.000 | 7.000 | 1.000 | 3.000 |
| 3. McFadden's R2 | 0.700 | 0.635 | 0.705 | 0.652 | 0.583 | 0.486 | 0.553 |
| 4. Null deviance | 52.192 | 52.192 | 52.192 | 104.385 | 104.385 | 104.385 | 156.577 |
| 5. Residual deviance | 15.659 | 19.036 | 15.380 | 36.348 | 43.570 | 53.635 | 69.916 |
| 6. AIC | 29.659 | 37.036 | 37.380 | 64.348 | 73.570 | 71.635 | 69.916 |
| 7. p-value of the Chi-sq statistic associated with the null deviance model | 0.186 | 0.186 | 0.186 | 0.128 | 0.128 | 0.128 | 0.089 |
| 8. p-value of the Chi-sq statistic associated with the residual deviance model | 0.999 | 0.990 | 0.997 | 0.999 | 0.998 | 0.991 | 0.999 |
| 9. p-value that determine whether there is significant reduction in deviance from null to residual | 2.62E-05 | 5.77E-05 | 6.10E-05 | 1.84E-09 | 8.44E-08 | 2.93E-08 | 4.14E-12 |
| 10. Hosmer and Lemeshow goodness of fit (GOF) test | 0.999 | 0.895 | 0.906 | 0.469 | 0.920 | 0.095 | 0.654 |

The Hosmer and Lemeshow goodness of fit test showed no significant difference between the logistic model and the observed data (p > 0.05) indicating that the logistic model constructed is a good fit (Table 1). The predicted probabilities are given in Table 2. Within the Order Artiodactyla, all species except *Bison bison bison* (American bison), *Ovis aries* (Sheep) and *Sus scrofa* (Pig) had more than 80% probability of viral (SARS-CoV-2) entry using ACE2 as a receptor. In American bison, Sheep and Pig, the probability of virus entry was 0.0036%, 24.3% and 18.6%, respectively. In Perrisodactyla, the probability of viral entry was 48% in horse and 79.1% in donkey. All the Carnivores in the study had a high probability of viral entry. In bats, the probability of viral entry was high in all the species. Amongst the rodents, except for Hamster, mouse and rat had a low probability of virus entry. The lagomorphs - rabbits and American pika had more than 90% probability of viral entry. All the primates had close to 100% probability of viral entry. The reptiles - Testudines and Crocodilia, showed medium to high probability of viral entry. However, in bird’s probability of viral entry varied, with chicken, golden eagle and duck having a low probability; and white-tailed eagles and turkey having a probability of 73.8% and 81%, respectively. Further, pangolins had a very high probability and African elephants a very low probability.

**Table 2.** Probability of viral entry in different species

| Class | Order | Family | Species (Common name) | Probability of Viral Entry (95% Confidence Interval) |
| --- | --- | --- | --- | --- |
| Mammalia | Artiodactyla | Bovidae | *Bos indicus* (Indian Cattle) | 9.98E-01(9.95E-01 – 1.00E+00) |
|  |  |  | *Bos taurus* (Exotic Cattle) | 9.17E-01(8.53E-01 – 9.55E-01) |
|  |  |  | *Bubalus bubalis* (Buffalo) | 8.25E-01(7.20E-01 – 8.96E-01) |
|  |  |  | *Bison bison bison* (American bison) | 3.60E-04(6.09E-05– 2.13E-03) |
|  |  |  | *Bos indicus* x *Bos taurus* (Indian crossbred Cattle) | 1.00E+00 (1.00E+00– 1.00E+00) |
|  |  | Camelidae | *Camelus bactrianus* (Double humped Camel) | 9.58E-01(9.19E-01 – 9.79E-01) |
|  |  |  | *Camelus dromedaries* (Single humped camel) | 9.58E-01(9.19E-01 – 9.79E-01) |
|  |  | Caprinae | *Capra hircus* (Goat) | 8.08E-01(7.06E-01 –8.80E-01) |
|  |  |  | *Ovis aries* (Sheep) | 2.43E-01(1.26E-01 –4.16E-01) |
|  |  | Suidae | *Sus scrofa* (Pig) | 1.86E-01(1.08E-01 – 3.02E-01) |
|  | Perissodactyla | Equidae | *Equus asinus* (Donkey) | 7.91E-01(6.77E-01 – 8.73E-01) |
|  |  |  | *Equus caballus* (Horse) | 4.80E-01(3.78E-01 – 5.85E-01) |
|  | Carnivora | Mustelidae | *Mustela putorius furo* (Ferret) | 9.99E-01(9.98E-01 – 1.00E+00) |
|  |  |  | *Lontra canadensis* (North American river otter) | 9.87E-01(9.71E-01 – 9.94E-01) |
|  |  | Felidae | *Panthera tigris altaica* (Siberian Tiger) | 8.92E-01(8.36E-01 – 9.31E-01) |
|  |  | Canidae | *Vulpes vulpes* (Red Fox) | 8.36E-01(7.71E-01 – 8.86E-01) |
|  |  |  | *Canis lupus familiaris* (Dog) | 9.78E-01(9.57E-01 – 9.88E-01) |
|  |  | Felidae | *Felis catus* (Cat) | 9.87E-01(9.71E-01 – 9.94E-01) |
|  | Chiroptera | Rhinolophidae | *Rhinolophus ferrumequinum* (Greater horseshoe bat) | 9.83E-01(7.71E-01 – 8.86E-01) |
|  |  | Phyllostomidae | *Desmodus rotundus* (Common vampire bat) | 9.88E-01(9.74E-01 – 9.94E-01) |
|  |  |  | *Phyllostomus discolor* (Pale spear-nosed bat) | 6.65E-01(5.49E-01 –7.64E-01) |
|  |  | Vespertilionidae | *Eptesicus fuscus* (Big brown bat) | 8.61E-01(7.82E-01 – 9.15E-01) |
|  |  |  | *Myotis brandtii* (Brandt's bat) | 9.12E-01(8.48 E-01 –9.51E-01) |
|  |  | Pteropodidae | *Pteropus Alecto* (Black fruit bat) | 9.98E-01(9.93E-01 – 9.99E-01) |
|  |  |  | *Rousettus aegyptiacus* (Egyptian fruit bat) | 1.00E+00 (9.99E-01 – 1.00E+00) |
|  | Rodentia | Cricetidae | *Cricetulus griseus* (Hamster) | 9.82E-01(9.59E-01 – 9.92E-01) |
|  |  | Muridae | *Mus musculus* (Mouse) | 4.97E-02(2.03E-02 – 1.17E-01) |
|  |  |  | *Rattus norvegicus* (Rat) | 2.87E-01(2.00E-01 – 3.94E-01) |
|  | Lagomorpha | Leporidae | *Oryctolagus cuniculus* (Rabbit) | 9.94E-01(9.86E-01 – 9.98E-01) |
|  |  | Ochotonidae | *Ochotona princeps* (American pika) | 9.66E-01(9.38E-01 – 9.81E-01) |
|  | Pholidota | Manidae | *Manis javanica* (Sunda pangolin) | 1.000E+00(1.00E+00–1.00E+00) |
|  | Primates | Hominidae | *Homo sapiens* (Human) | 1.00E+00(9.99E-01 – 1.00E+00) |
|  |  | Cercopithecoidea | *Macaca fascicularis* (Crab eating monkey) | 1.00E+00(1.00E+00 – 1.00E+00) |
|  |  |  | *Macaca mulatta* (Rhesus monkey) | 1.00E+00(9.99E-01 – 1.00E+00) |
|  |  |  | *Macaca nemestrina* (Southern pig-tailed monkey) | 1.00E+00(1.00E+00 – 1.00E+00) |
|  |  | Hominidae | *Pan troglodytes* (Chimpanzee) | 9.99E-01(9.98E-01 - 1.00E+00) |
|  |  | Cercopithecidae | *Papio Anubis* (Baboon) | 1.00E+00(9.99E-01 – 1.00E+00) |
|  | Proboscidea | Elephantidae | *Loxodonta Africana* (African elephant) | 2.08E-01(1.40E-01 – 2.99E-01) |
| Reptiles | Testudines | Cheloniidae | *Chelonia mydas* (Green sea turtle) | 7.71E-01(7.06E-01 – 8.26E-01) |
|  |  | Emydidae | *Chrysemys picta bellii* (Painted turtle) | 4.96E-01(3.55E-01 – 6.39E-01) |
|  |  | Trionychidae | *Pelodiscus sinensis* (Chinese softshell turtle) | 5.92E-01(4.03E-01 – 7.57E-01) |
|  | Crocodilia | Alligatoridae | *Alligator sinensis* (Chinese alligator) | 9.93E-01(9.80E-01-9.98E-01) |
|  |  | Crocodylidae | *Crocodylus porosus* (Saltwater alligator) | 9.82E-01(9.55E-01 – 9.93E-01) |
| Aves | Galliformes | Phasianidae | *Gallus gallus* (Chicken) | 4.84E-03(1.59E-03 – 1.46E-02) |
|  |  |  | *Meleagris gallopavo* (Turkey) | 8.15E-01(6.86E-01 – 8.99E-01) |
|  | Anseriformes | Anatidae | *Anas platyrhynchos* (Mallard) | 1.91E-03(4.31E-04 – 8.46E-03) |
|  | Accipitriformes | Accipitridae | *Haliaeetus albicilla* (White-tailed eagle) | 7.38E-01(5.96E-01 – 8.42E-01) |
|  |  |  | *Aquila chrysaetos chrysaetos* (Golden Eagle) | 3.32E-02(1.54E-02– 7.02E-02) |

## Discussion

Recognition of the receptor is an important determinant in identifying the host range and cross-species infection of viruses[24]. It has been established that ACE2 is the cellular receptor of SARS-CoV-2[16]. This study is targeted to predict viral entry in a host, *i.e.,* hosts that can be reservoir hosts (Artiodactyla, Perrisodactyla, Chiroptera, Carnivora, Lagomorpha, Primates, Pholidota, Proboscidea, Testudines, Crocodilia, Accipitriformes and Galliformes) and hosts that can be appropriate small animal laboratory models (Rodentia) of SARS-CoV-2, through sequence comparison, homology modeling of ACE2, docking the modelled homology structures with the spike - ACE2 binding domain and prediction of viral entry.

Initially for prediction of probability of viral entry, sequence comparison of ACE2 was done vis - a - vis, within group distance; distance of an Order from the Order primates, distance of each individual taxa from humans; variability in the ACE2 spike interacting domain at protein and nucleotide level; and phylogeny. Considering the pandemic nature of the disease in humans, the low within-group distance in primates indicated that all the species considered within the Order primates are prone to be equally infected with SARS-CoV-2 as humans. On comparing the Orders, Galliformes was most distant from the primates and carnivora was found proximal. This confirms to the recent reports of chicken (Galliformes) and ducks (Anseriformes) not being infected with SARS-CoV-2 [22], and tigers and lions being infected[12]. On comparing individual hosts, pig was found to be the established taxa that is uninfected with SARS-CoV-2 [22]. Considering the distance of pig from *Homo sapiens* as a cut-off, would include all the carnivores, perissodactyls and few artiodactyls viz. goat, buffalo, bison and sheep, to be infected, but, excludes cattle (Artiodactyla), all bats (Chiroptera) and birds (Galliformes, Anseriformes and Accipitriformes). Further, the negative selection observed on codon-based test of neutrality, indicates that, the variation at the nucleotide level, is translated synonymously, indicating that the structure of ACE2 is conserved through the process of evolution. The comparison of the spike binding domains across all the Orders, also did not lead to meaningful conclusions on viral entry in different species,

On phylogeny, sub-clustering of the rodents, lagomorphs and carnivores close to primates with reliable bootstrap values partially corroborates with the occurrence of SARS-CoV-2 infection in carnivores [22] as mice were found not infected with SARS-CoV-2 [16]. Further, sub-clustering of fruit-bat with horseshoe bat suggests possible entry of the virus in fruit-bat, as the COVID-19 outbreak in Wuhan in Dec 2019 was traced back to have a probable origin from horseshoe bat [16]. The virus strain RaTG13 isolated from this bat was found to have 96.2% sequence similarity with the human SARS-CoV-2. These results again led to no concrete conclusions on viral entry in various hosts. Therefore, to assess the probability of viral entry in various hosts, after homology modeling of ACE2 and docking the modelled homology structures with the spike – ACE2 binding domain, 32 spike binding parameters were evaluated.

A total of 9 data for each host for each spike binding parameter as described in the materials and methods are available to select the parameters that would clearly classify the Orders into infected/uninfected. However, none of the 6 parameters – RMSD, delta G, Intraclashes Group1, Van der Waals, Solvation Hydrophobic and entropy sidechain, that were significantly different in the infected from the uninfected could classify the Orders into infected or uninfected. This suggests that a single parameter at a time, as has been considered in recent reports[21], may not be considered and evaluated for estimating the probability of virus entry. Therefore, logistic regression with all the estimated parameters was done with seven different combination of data to predict the probability of viral entry. The best combination of X-ray crystallography models was identified based on evaluation parameters – Number of parameters significant in the model at 1% LS, Number of parameters significant in the model at 5% LS, McFadden R^2^, Null deviance, Residual deviance, AIC, p-value of the Chi-sq statistic associated with the null deviance model, p-value of the Chi-sq statistic associated with the residual deviance model, p-value to determine whether there is significant reduction in deviance from null to residual and Hosmer and Lemeshow goodness of fit (GOF) test.

McFadden R^2^ is a measure of fit in statistical modeling [31]. However, this can be used only to compare models with same number of covariates i.e. this increase with an additional covariate. Akaike information criterion (AIC) is used to compare models fitted over same datasets. Lower the AIC better is the model and better is the fit [32], Significant reduction in the null deviance is assessed by the change in the p-value of the Chi-sq statistic associated with the null deviance model to the p-value of the Chi-sq statistic associated with the residual deviance model. This can be further determined by the p-value that determines whether there is significant reduction in deviance from null to residual. A non-significant p-value on Hosmer and Lemeshow goodness of fit (GOF) test indicates that there is no evidence that the model is not fitting well with the data considered. All these parameters were relatively better for the data against the combinations - 6LZG & 6M0J; 6LZG & 6VW1 and 6M0J & 6VW1 & 6LZG than the other four combinations. The number of significant parameters at 1% and 5% level of significance were greater in these combinations than the other four. The reduction in null deviance was found to be highly significant in 6M0J & 6VW1 & 6LZG followed by 6LZG & 6M0J and 6LZG & 6VW1. Considering several criteria as mentioned, the data against these models were finally considered to predict the probability of viral entry on the test data and the prediction accuracy was found to be higher for the data against 6LZG & 6M0J.

Root-Mean-Square-Distance (RMSD) was the most significant parameter amongst the 32 spike binding parameters of ACE2 in all the logistic models considered (Supplementary File 1). RMSD measures the degree of similarity between two optimally superposed protein 3D structures [33]. The smaller the RMSD between two structures, more similar they are. Docking predictions within an RMSD of 2 Å are considered successful, whereas values higher than 3 Å indicate docking failures [34]. The average RMSD in the infected and uninfected known hosts was 0.068 and 0.113, respectively. In all the logistic models, the coefficient (i.e. the log of odds ratio) of RMSD was negative, indicating that RMSD is negatively connected with infection. This means that the increase in RMSD would lead to higher odds of not getting infected. In the combination that is finalized (i.e. combination of 6LZG & 6M0J) for predicting the probability of viral entry, the coefficient of RMSD was −5.575e+01. Further, the deviance residuals for this logistic model from this combination were symmetric as indicated by median (0.01172), which is close to zero. The AIC for this selected combination is 64.348. Further, there was also a significant reduction in null deviance with an R-square of 0.652. The prediction equation on analysis of these data against the combination 6LZG & 6M0J, was used to predict the probability of viral entry in various hosts.

As observed in this study, it has been predicted that *Bos indicus* (Indian cattle) and *Bos taurus* (Exotic cattle) can act as intermediate hosts of SARS-CoV-2 [27] and that pigs are not susceptible [22]. Also, Camels, which are reported to be infected with SARS-CoV [28] are equally capable of SARS-CoV-2 infection. Among the rodents, hamsters had the highest probability of viral entry. It has been established that SARS-CoV-2 effectively infects hamster[29] and, rats and mice were found less probable[26]. All the Carnivores in the study had high probability of viral entry. Reports of SARS-CoV2 infection in cats[22], tigers and lions[12] substantiate our estimates obtained in the study. Rabbits also had high probability of viral entry showing concordance to the recent evidence of SARS-CoV-2 replication in rabbit cell lines[30]. All the primates close to human species were identified to be highly probable. The variability within the Order(s) must be reason for not being able to classify them as a group, to either being infected or uninfected using unpaired t-test.

## Conclusion

Most of the species considered under different Orders, in this study, showed high probability of viral entry. The findings hint towards the probable hosts that can act as laboratory models or as reservoir hosts and allows us to take a cue about the probable pathogenic insult that can be caused by SARS-CoV-2 to different species. This, however, warrants further research. Also, viral entry is not the only factor that determines infection in COVID-19 as viral loads were found to be high in asymptomatic patients [35, 36]. The important factors that determine disease/infection(COVID-19) in host(s) are - Host defense potential, underlying health conditions, host behavior and number of contacts, Age, Atmospheric temperature, Population density, Airflow and ventilation and Humidity[37].

## Materials and methods

Sequence analysis, phylogenetic analysis, homology modeling of ACE2, docking the modelled homology structures with the spike - ACE2 binding domain and prediction of viral entry were done in this study (Figure 5).

**Figure 5.**
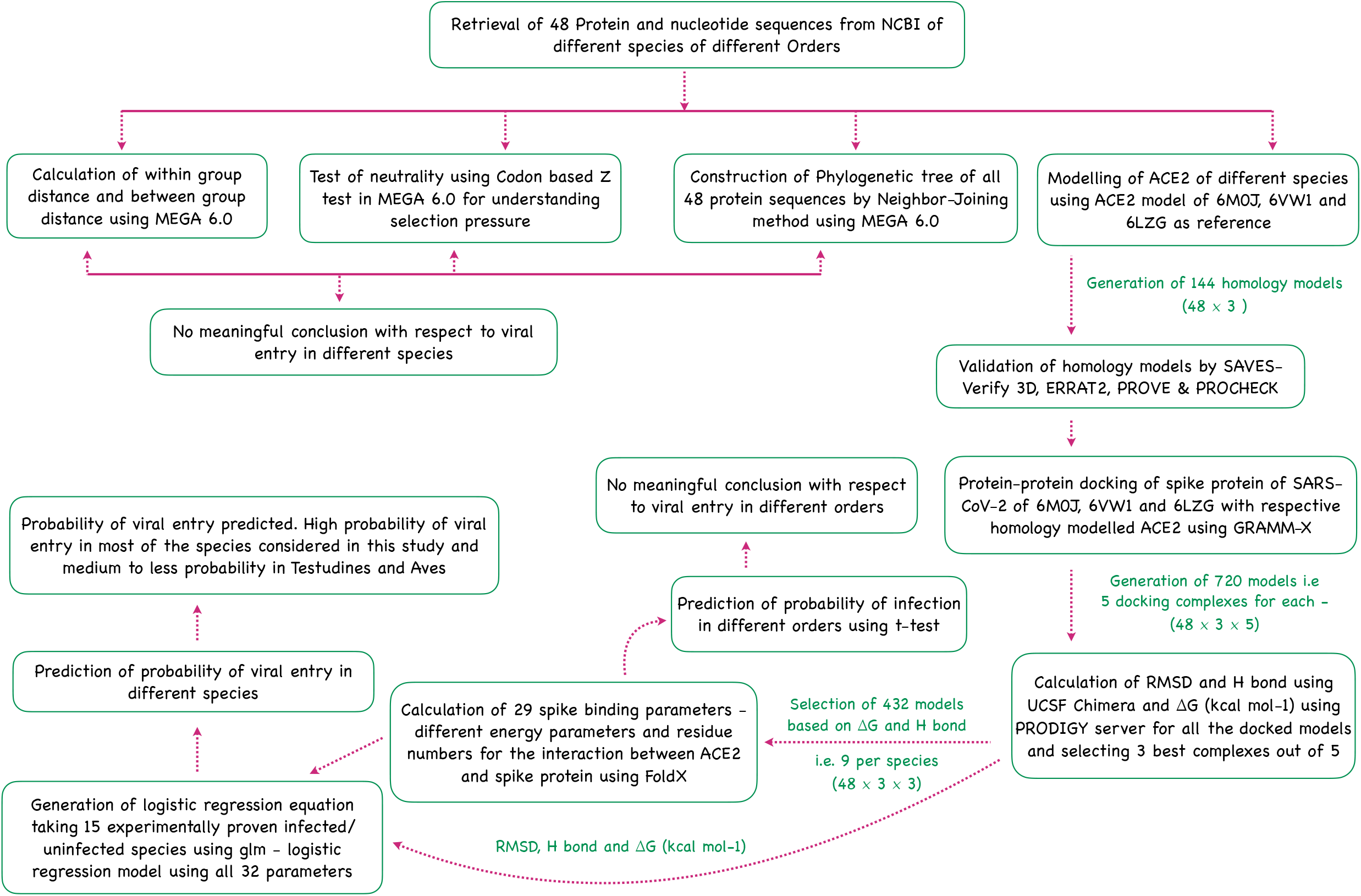
Flowchart showing the step wise analysis for the work carried out to estimate the probability of virus entry.

### Sequence analysis

In this study, 48 (mammalian, reptilian and avian species) ACE2 complete/partial protein and nucleotide sequences available on NCBI were analyzed (Supplementary Table 1) to understand the possible difference(s) in the ACE2 sequences that may correlate with SARS-CoV-2 viral entry into the cell. The partial sequences are considered in the study after ensuring that these sequences completely cover the spike interacting domain of ACE2. Within the mammalian class, Orders - Artiodactyla, Perrisodactyla, Chiroptera, Rodentia, Carnivora, Lagomorpha, Primates, Pholidota and Proboscidea; within the Reptilian class, Orders - Testudines and Crocodilia; and within the Avian class, Orders - Accipitriformes, Anseriformes and Galliformes, were considered in the study. These Orders were considered keeping in view all the possible reservoir hosts/ laboratory animal models that can possibly be infected with the SARS-CoV-2. The within and between group distances were calculated in Mega 6.0[38]. The ACE2 sequences in the study, are compared as a group (average of the Order) with the average of all species in the Order Primates or individually with the *Homo sapiens*ACE2 sequence. The Codon-based Z test of selection (strict-neutrality (dN=dS)) to evaluate synonymous and non-synonymous substitutions across the ACE2 sequences among the Orders was done. Further, for comparing the sequence of the spike interacting domain, this was identified to be defined in the UniProt ID - Q9BYF1. The family and domains section of the UniProt ID Q9BYF1 clearly marks the sequence location of the ACE2 - spike interacting domains as 30 - 41aa, 82 - 84 aa and 353 - 357 aa. The nucleotide sequence alignments at positions that correspond to the spike-binding domain of *Homo sapiens* ACE2 are 90-123 bp; 244-252 bp and 1058-1071 bp.

### Phylogenetic analysis

Phylogenetic analysis of the protein sequences was done using MEGA 6.0[38]. Initially, the sequence alignment was done using Clustal W[39]. The aligned sequences were then analyzed for the best nucleotide substitution model on the basis of Bayesian information criterion scores using the JModelTest software v2.1.7[40]. The tree was constructed by the Neighbor-joining method with the best model obtained using 1000 bootstrap replicates. It is important to note that the missing data or gaps are treated in this analysis by using pair-wise deletion.

### Homology modeling

The Structures of novel coronavirus spike receptor-binding domain complexed with its receptor - ACE2, that were determined through X-ray diffraction are available at PDB database with IDs 6LZG [25], 6M0J [41] and 6VW1 [42]These available ACE2 models from PDB database were used for homology modeling using SWISS-MODEL[43], which was accessed through ExPASy web server. The models (144 = 48 × 3) were validated through SAVES [44]. SAVES is a conglomerate of different validating algorithms like PROCHECK, VERIFY 3D, ERRAT2, PROVE. The models are assigned “PASS’ by Verify 3D when more than 80% of the amino acids have scored ≥ 0.2 in 3D/1D profile. In case of ERRAT2, models thar scored more than 95% are considered to have good resolution. PROVE gives: Error (>5%), Warning (1 to 5%) or Pass (<1%) based on% of buried atoms. From PROCHECK, Ramachandran plot with over 90% of the residues in core regions is considered to be a good model.

### Protein-protein Docking

The spike ACE2 - binding domains of 6LZG, 6M0J and 6VW1 were used in docking along with the respective homology modelled structures of ACE2 protein of all the hosts, *i.e.*, ACE2 of 48 hosts as a receptor and spike ACE2 binding domain of SARS-CoV-2 as a ligand for protein-protein docking. GRAMM-X docking server was used for protein-protein docking, which generated a docked complex [45]. Five docked complexes were generated from GRAMM-X for each X-ray crystallography model in each species and post-docking analyses was carried out using Chimera software[46] and PRODIGY [47]. A total of 720 models (48 hosts × 3 X-ray Crystallography models × 5 docking complexes) were analyzed. Chimera is an extensible program for interactive visualization and analysis of molecular structures for use in structural biology. Chimera provides the user with high quality 3D images, density maps, trajectories of small molecules and biological macromolecules, such as proteins. The number of hydrogen bonds in each docking structure was estimated using Chimera and the delta G of the docked models was estimated using PRODIGY.

Out of the five docked complexes generated through GRAMM-X, three best complexes for each host under each X-Crystallography structure were selected (432 model = 48 × 3 × 3) for further analysis based on delta G and number of hydrogen bonds (Supplementary Figure 3 and Supplementary Table 6). The docked models are expected to differ from the real structure and the differences are quantified by root mean square deviation (RMSD).To estimate RMSD (root mean squared deviation) the three best docked complexes of each X-ray crystallography model in each species were compared with the respective models-6LZG/6M0J/6VW1 using Chimera. Further, in addition to delta G and RMSD, in FoldX software [48] several parameters were estimated for all these selected docked structures (for 432 models (48 hosts × 3 X-ray Crystallography models × 3 selected docking complexes) were analyzed), These parameters include - IntraclashesGroup1, IntraclashesGroup2, Interaction Energy, Backbone Hbond, Sidechain Hbond, Van der Waals, Electrostatics, Solvation Polar, Solvation Hydrophobic, Van der Waals clashes, entropy sidechain, entropy mainchain, sloop entropy, mloop entropy, cis bond, torsional clash, backbone clash, helix dipole, water bridge, disulfide, electrostatic kon, partial covalent bonds, energy Ionisation, Entropy complex, Number of Residues, Interface Residues, Interface Residues Clashing, Interface Residues VdW Clashing and Interface Residues BB Clashing. All these 32 parameters (29 in FoldX, delta G, H bonds and RMSD) are referred to as spike binding parameters of ACE2.

### Statistical analysis for prediction

Till date, clear-cut information of 15 species that are either infected or uninfected with SARS-CoV2 is available (Supplementary Table 7). For each of these species, a total of nine models with their parameters were taken for the analysis i.e. for each species, the three selected docked structures for each of the X-ray crystallography structures were selected (Supplementary Figure 3). A total of 135 data per parameter (15 hosts × 3 X-ray Crystallography models × 3 selected docking complexes) were analyzed. Initially, for each parameter (spike binding parameters of ACE2), the difference between the infected and uninfected is evaluated using Unpaired t-test in GraphPad Prism 7.00 (GraphPad Software, La Jolla, California, USA). Welch correction was applied wherever necessary. For those parameters that were significant, the difference between Order(s) means and the infected/uninfected groups was also further evaluated using Unpaired t-test(Note: if a species is included in the infected/uninfected group, the same is not included in its Order on comparing the Order(s) with infected/uninfected group) (Supplementary table 9 for more information).

Later, backward stepwise logistic regression model was constructed on all the 32 parameters (29 from FoldX, RMSD, H bonds and delta G) estimated above in the 15 known species of infected (11) and uninfected (4) (Supplementary Table 7). A total of 135 data per parameter were available across the three X-ray Crystallography structures considered. These data were used in seven different combinations based on the combination of X-ray Crystallography structures. The seven combinations include, data against single model - 6LZG,6M0J and 6VW1 (i.e. 45 data); data against two models - 6LZG and 6M0J / 6LZG and 6VW1 / 6M0J and 6VW1 (i.e. 90 data); and data against all the three models - 6LZG and 6M0J and 6VW1 (i.e. 135 data). These seven combinations were evaluated based on the estimates of Number of parameters significant in the logistic model at 1% LS, Number of parameters significant in the logistic model at 5% LS, McFadden’s R2, Null deviance, Residual deviance, AIC, p-value of the Chi-sq statistic associated with the null deviance model, p-value of the Chi-sq statistic associated with the residual deviance model, p-value to determine whether there is significant reduction in deviance from null to residual, Hosmer and Lemeshow Goodness of fit (GOF) test. After selecting the best combination(s), the best model (prediction equation) was selected after evaluation of the training and test data sets for each of the combinations. This prediction equation from the best combination of data was used to predict the probability of viral entry in rest of the species using the average values of the top three models for all the parameters in the equation.

Further, with 32 parameters, the minimum sample size required to derive statistics that represent each parameter, is 1700[50] *(n =100 + xi i.e. here: −n = 100 + (100 + (50 × 26) = 1700, with a minimum of 50 events per parameter).* The data was needed to be extrapolated to at least 1700 to predict the confidence intervals. This was based on the assumption that the ACE2 structure and sequence is conserved within a species. For the species -*Homo sapiens*, we compared several ACE2 sequences and found that all the compared sequences were identical. With this assumption that the spike binding parameters of ACE2 within a species are conserved and due to the pandemic nature of the disease the data was extrapolated.

## Abbreviations

ACE2: Angiotensin-converting enzyme 2
CDS: Coding Sequence
COVID-19: Coronavirus disease 2019
ICTV: International Committee on Taxonomy of Viruses
MERS: Middle East Respiratory Syndrome
PDB: Protein Data Bank
RMSD: Root-mean-square deviation
SADS: Swine Acute Diarrhea Syndrome
SARS-CoV-2: Severe Acute Respiratory Syndrome Coronavirus 2
SARS-CoV: Severe Acute Respiratory Syndrome Coronavirus
SARS: Severe Acute Respiratory Syndrome
WHO: World Health Organization

---

**Supplementary Figure 1.** Nucleotide sequence alignment of the CDS region of ACE2. The shaded regions show the spike interacting domains.

**Supplementary Figure 2.** Protein sequence alignment of ACE2. The shaded regions show the spike interacting domains.

**Supplementary Figure 3.** Depiction of numbers of models considered in this study showing the number of values per parameter. For each species the ACE2 sequence is homology modelled against the three X-crystallography structures - 6M0J,6LZG and 6VW1. The spike ACE2 binding domain of each of the X-crystallography structures is docked with its homology modelled ACE2 and 5 docked complexes were evaluated to select the top three models. This leaves us with 9 values for all the spike binding parameters for further analysis.

**Supplementary Table 1.** Species considered in this study

**Supplementary Table 2**. Within Mean group distance among the Orders

**Supplementary Table 3**. Between group distance (between Primates and other groups)

**Supplementary Table 4.** Evaluation of homology modelled structures through SAVES. ACE2 sequence of each species is homology modelled against the three X-crystallography structures - 6M0J,6LZG and 6VW1. This excel file contains three sheets, each sheet is for each of the three X-crystallography structures. A total of 144 homology modelled structures were evaluated (48 for each three X-crystallography structures)

**Supplementary Table 5**. Parameters obtained from UCSF Chimera and PRODIGY for 720 models. For each species the ACE2 sequence is homology modelled against the three X-crystallography structures - 6M0J,6LZG and 6VW1. The spike ACE2 binding domain of each of the X-crystallography structures is docked with its homology modelled ACE2 and 5 docked complexes were evaluated. This leaves us with 720 models (48 × 3 × 5) to be evaluated using delta G and H bonds.

**Supplementary Table 6.** Parameters obtained from FoldX for the 432 models. For each species the ACE2 sequence is homology modelled against the three X-crystallography structures - 6M0J,6LZG and 6VW1. The spike ACE2 binding domain of each of the X-crystallography structures is docked with its homology modelled ACE2 and 5 docked complexes were evaluated to select the top three models. This leaves us with 432 models (48 × 3 × 3) for the final analysis.

**Supplementary Table 7.** Lists of experimentally proven infected/uninfected (Infected-1 and Uninfected-0) animals with other spike binding parameters. A total of 135 data per parameter (15 hosts × 3 X-ray Crystallography models × 3 selected docking complexes) were considered for logistic model construction

**Supplementary Table 8.** List of significant spike binding parameters after Unpaired t-test between the known infected and uninfected groups

**Supplementary Table 9.** Data considered for evaluating the Order from the uninfected and infected groups by unpaired t-test

**Supplementary File 1.** Details about the commands used and results obtained after testing different combination of models.

## Author’s contributions

MRP performed sequence alignment and phylogeny of nucleotide and amino acid and drafted the manuscript. PG, SS, VK & NT performed protein modelling and docking and estimated the different parameters from FoldX. RINK retrieved the amino acid and nucleotide sequences and edited the manuscripts. MP, GSK and BM edited and proofread the manuscript. RKG did complete statistical analysis and manuscript development. TM, SM, RKS, RKG and BPM conceptualized and planned the entire study.

## Acknowledgments

We are grateful to Director NIAB and Director IVRI for the support.

