## Supplementary figures and images for "SARS-CoV-2 Spike Glycoprotein and ACE2 interaction reveals modulation of viral entry in wild and domestic animals"

### Supplementary Figure 1

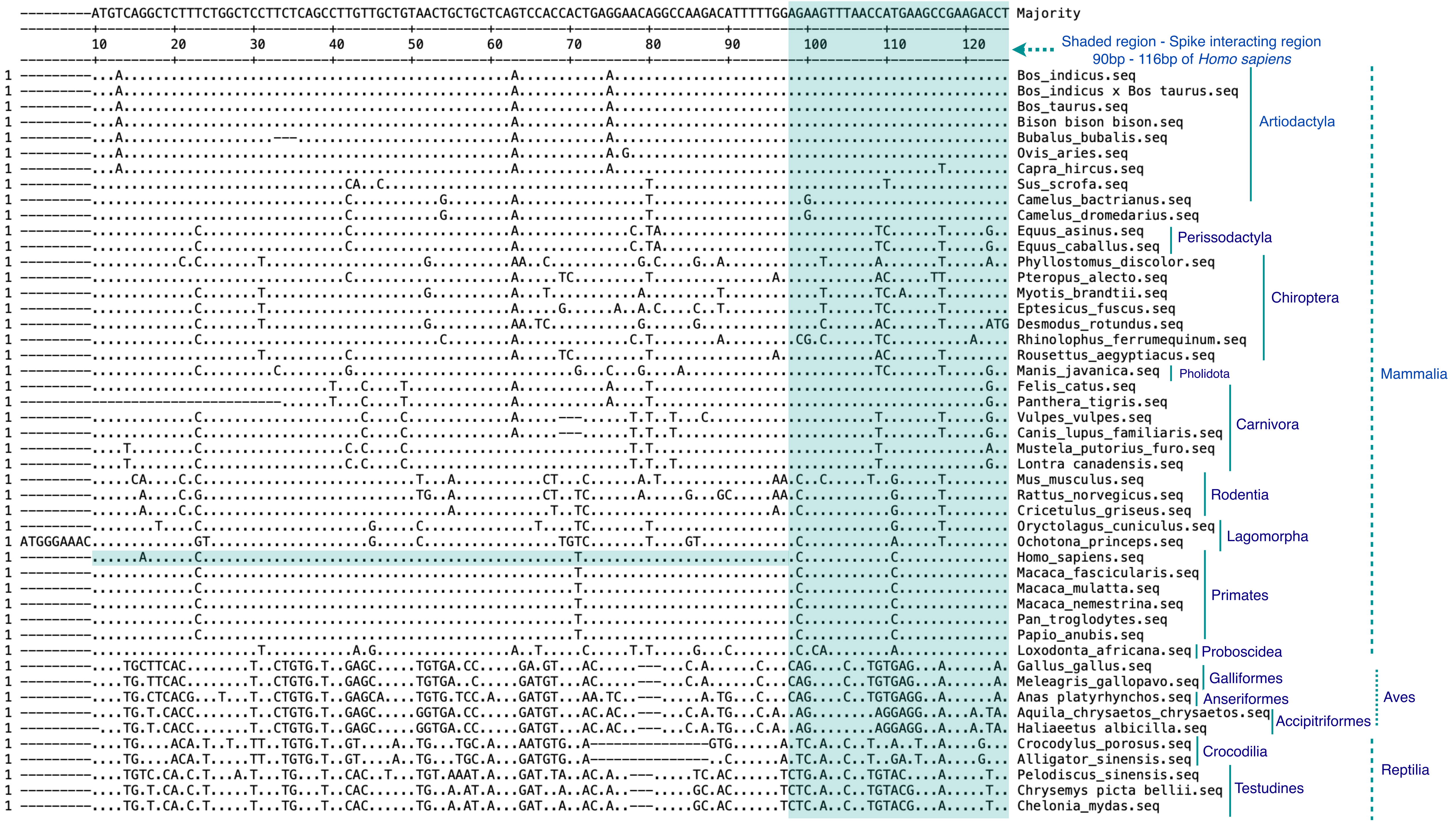

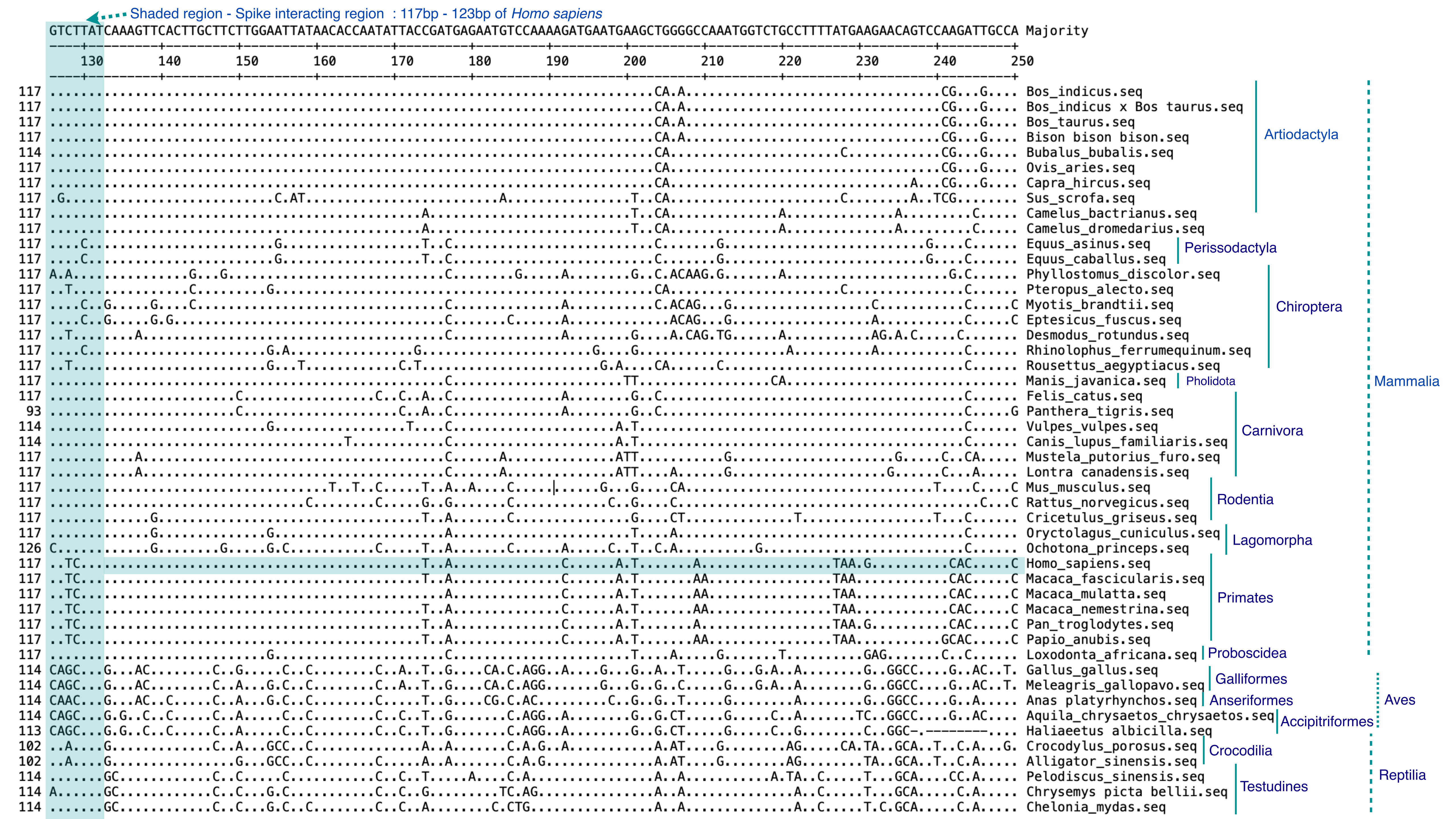

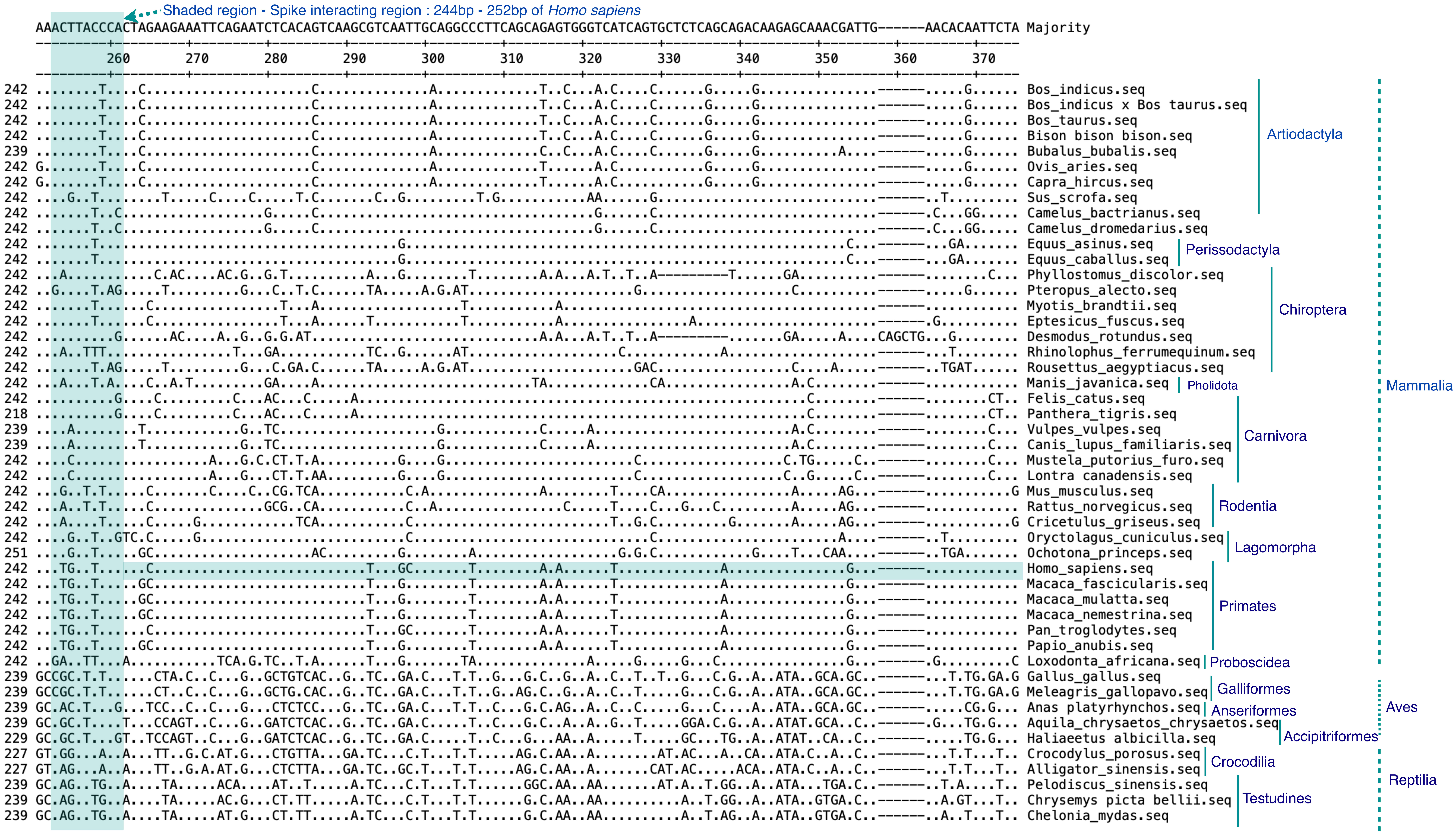

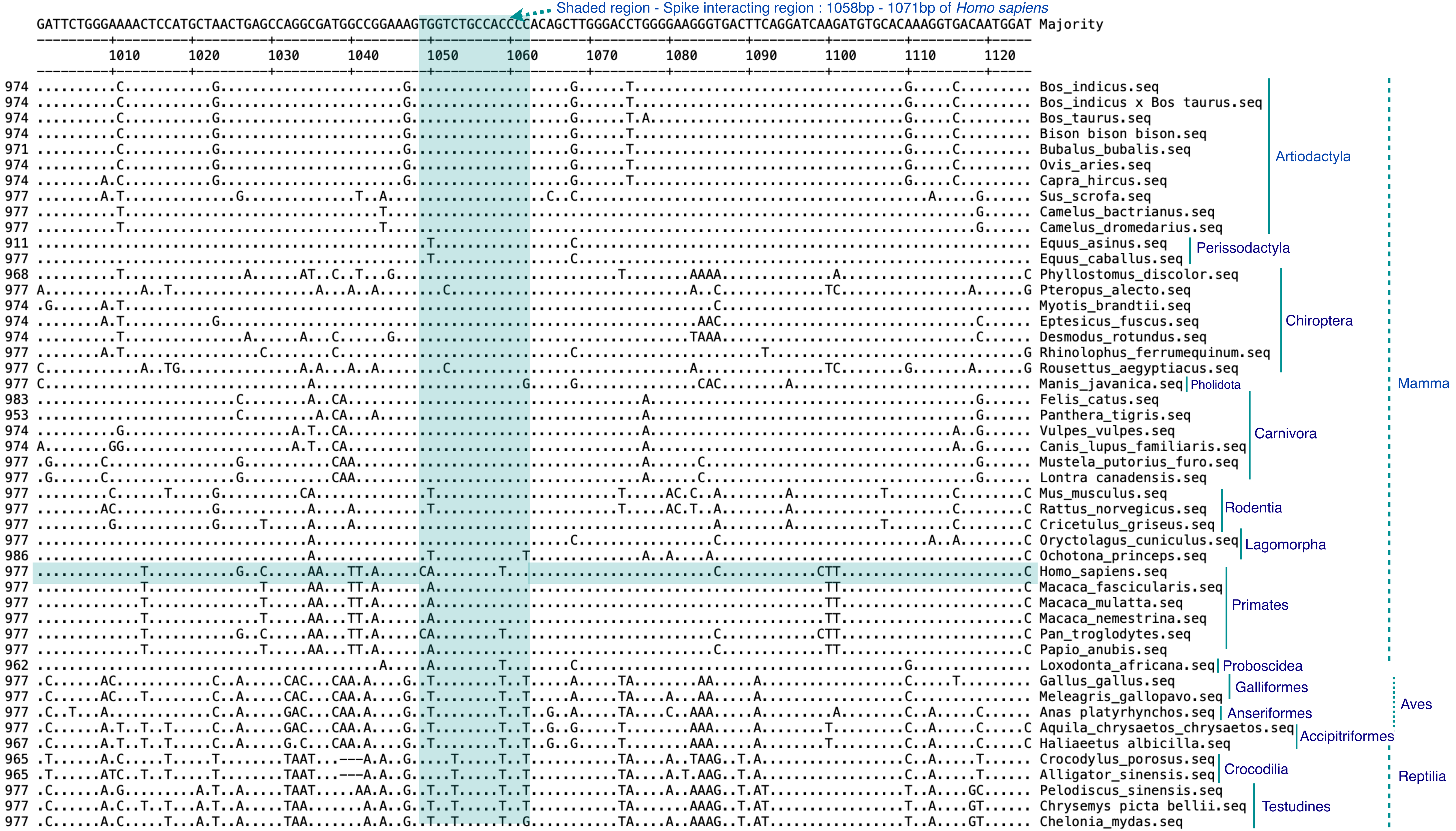

### Supplementary Figure 3

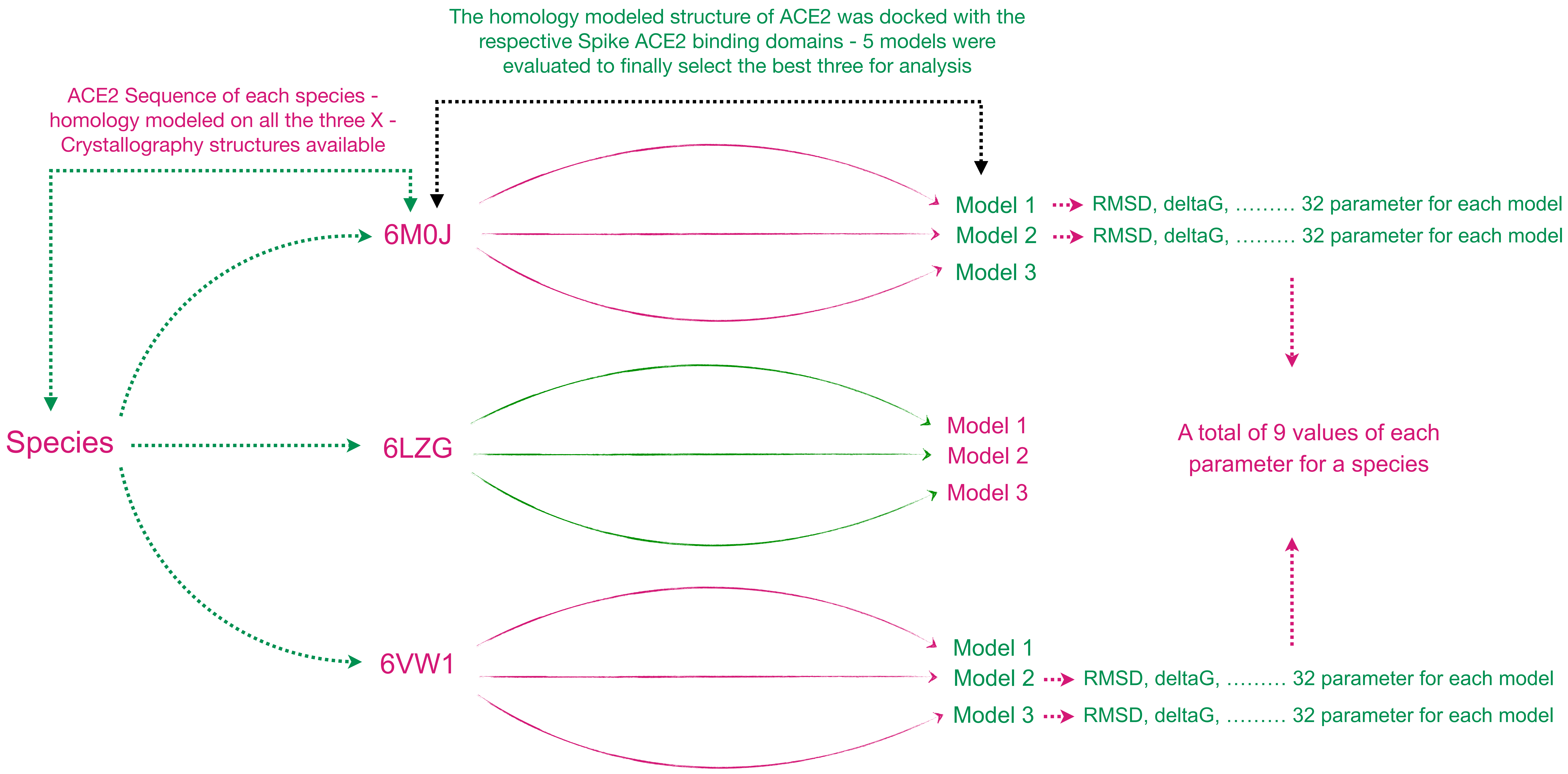
