## Supplementary Figure 2 for "SARS-CoV-2 Spike Glycoprotein and ACE2 interaction reveals modulation of viral entry in wild and domestic animals"

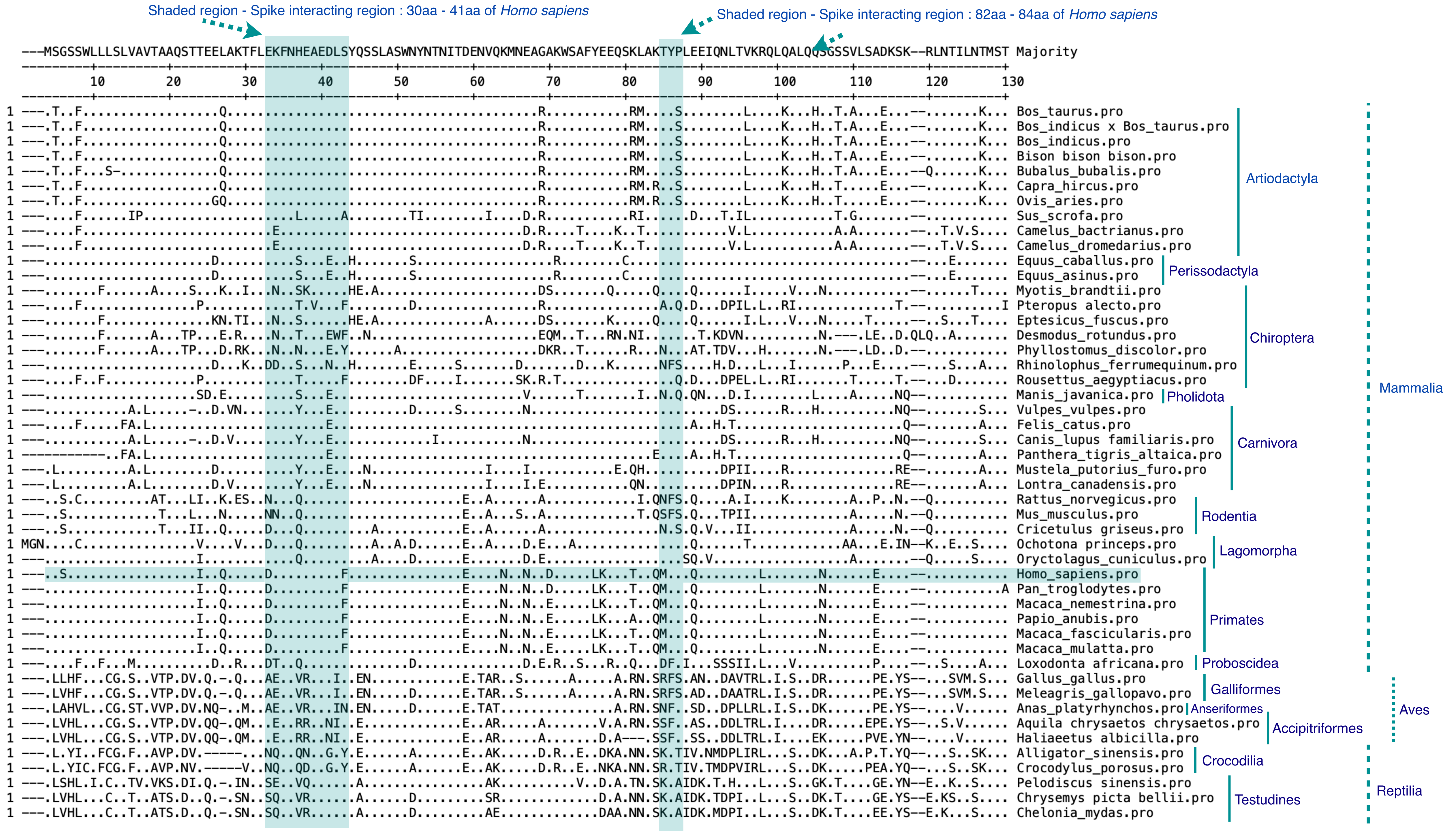

Shaded region - Spike interacting region : 30aa - 41aa of Homo sapiens
Shaded region - Spike interacting region : 82aa - 84aa of Homo sapiens
---MSGSSWLLLSLVAVTAAQSTTEELAKTFLEKFNHEAEDLSYQSSLASWNYNTNITDENVQKMNEAGAKWSAFYEEQSKLAKTYPLEEIQNLTVKRQLQALQQSGSSVLSADKSK--RLNTILNTMST Majority
Bos_taurus.pro
Bos_indicus x Bos_taurus.pro
Bos_indicus.pro
Bison bison bison.pro
Bubalus_bubalis.pro
Capra_hircus.pro
Ovis_aries.pro
Sus_scrofa.pro
Camelus_bactrianus.pro
Camelus_dromedarius.pro
Artiodactyla
Equus_caballus.pro
Equus_asinus.pro
Perissodactyla
Myotis_brandtii.pro
Pteropus alecto.pro
Eptesicus_fuscus.pro
Desmodus_rotundus.pro
Phyllostomus_discolor.pro
Rhinolophus_ferrumequinum.pro
Rousettus_aegyptiacus.pro
Chiroptera
Manis_javanica.pro
Pholidota
Vulpes_vulpes.pro
Felis_catus.pro
Canis_lupus familiaris.pro
Panthera_tigris_altaica.pro
Mustela_putorius_furo.pro
Lontra_canadensis.pro
Carnivora
Rattus_norvegicus.pro
Mus_musculus.pro
Cricetulus griseus.pro
Rodentia
Ochotona princeps.pro
Oryctolagus_cuniculus.pro
Lagomorpha
Homo_sapiens.pro
Pan_troglodytes.pro
Macaca_nemestrina.pro
Papio_anubis.pro
Macaca_fascicularis.pro
Macaca_mulatta.pro
Primates
Loxodonta africana.pro
Proboscidea
Mammalia
Gallus_gallus.pro
Meleagris_gallopavo.pro
Galliformes
Anas_platyrhynchos.pro
Anseriformes
Aquila chrysaetos chrysaetos.pro
Haliaeetus albicilla.pro
Accipitriformes
Aves
Alligator_sinensis.pro
Crocodylus_porosus.pro
Crocodilia
Pelodiscus sinensis.pro
Chrysemys picta bellii.pro
Chelonia_mydas.pro
Testudines
Reptilia

Shaded region - Spike interacting region : 353aa - 357aa of *Homo sapiens*

```
    SY-ISPTGCLPAHLLGDMWGRFWTNLYSLTVPFGQKPNIDVTDAMVNQSWDAERIFKEAEKFFVSVGLPNMTQGFWENSMLTEPGDGRKVVCHPTAWDLGKGDFRIKMCTKVTMDDFLTAHHEMGHIQYD Majority
    ---------+---------+---------+---------+---------+---------+---------+---------+---------+---------+---------+---------+---------+
            270       280       290       300       310       320       330       340       350       360       370       380       390
    ---------+---------+---------+---------+---------+---------+---------+---------+---------+---------+---------+---------+---------+
253 ..-..............................EH..S....EK.E...................IS..Y......D..................................................... Bos_taurus.pro
253 ..-..............................EH..S....EK.E...................IS..Y......D..................................................... Bos_indicus x Bos_taurus.pro
253 ..-..............................EH..S....EK.E...................IS..Y......D..................................................... Bos_indicus.pro
253 ..-..............................EH..S....EK.E...................IS..Y......D..................................................... Bison bison bison.pro
252 ..-..............................EH..S....EK.E...................IS..Y......D..................................................... Bubalus_bubalis.pro   Artiodactyla
253 ..-..............................EH..S....EK.K...................I...Y......N..................................................... Capra_hircus.pro
253 ..-..............................EH..S....EK.K...................I...Y......D..................................................... Ovis_aries.pro
254 .R-........................P......E..S....E.........I...E........I..........N..................................................... Sus_scrofa.pro
254 .H-.......................................E..E......K............I..........D..................................................... Camelus_bactrianus.pro
254 .H-.......................................E..E......K............I..........D..................................................... Camelus_dromedarius.pro
254 .H-.N.........................................D.....K...E......................................................................... Equus_caballus.pro   Perissodactyla
240 D---TDV.------QC..............................D.....K...E......................................................................... Equus_asinus.pro
253 .R-.....Y..................N.....E........G...E......K........YI.....S..P...N..................................................... Myotis_brandtii.pro
254 .H-....................I...Q.....E.........E....N..EK............L......EK...K.......NDQ..A...............I.....K.E............... Pteropus alecto.pro
253 .R-........................N.....E...........KE......K........YM.....S..P...N........................N............................ Eptesicus_fuscus.pro
253 .H-........................N..A...E..T....A...D.....Q..........K....FS......D.....K.D...E...........NK............................ Desmodus_rotundus.pro   Chiroptera
251 .R-........................D..A..PE..T....S...A.....Q...............F.......D.....K.D...E............K............................ Phyllostomus_discolor.pro
254 FH-........................P.................L..N...K............I......E...N.....D...............................E............... Rhinolophus_ferrumequinum.pro
254 .H-....................I...P.....E.........E....N.N.K............L......ET...K.V....DNDQ..A...............I.....K.E..............Y Rousettus_aegyptiacus.pro
254 .H-........................P.....R..............T...N................K...T...........................H............................ Manis_javanica.pro   Pholidota   Mammalia
253 ..-........................P..............N.........RK..............................S.S........................................... Vulpes_vulpes.pro
256 .R-........................P........................R.................................S........................................... Felis_catus.pro
253 ..-........................P..............N.........RK...................E..G.......S.S........................................... Canis_lupus familiaris.pro   Carnivora
246 .R-........................P........................R................................NSQ.......................................... Panthera_tigris_altaica.pro
254 .R-........................P.M...R..................R...E...T...........E...Q.........N..............R............................ Mustela_putorius_furo.pro
254 .R-........................P.M..................R...R...E...............E...Q.........N..............R............................ Lontra_canadensis.pro
254 ..-........................P..T..L...................................Q..P...T.........D.............H..............N.............. Rattus_norvegicus.pro
254 ..-........................P.....A...........M..G.......Q............H......A.......A...............H..............N.............. Mus_musculus.pro   Rodentia
254 .F-........................P....................G....................H......G.....D...D............................N.............. Cricetulus griseus.pro
257 .Q-M.A.....................................T....G....................S...............................D.............NY............. Ochotona princeps.pro
254 .R-........................................T....G....................S.............................................N.............. Oryctolagus_cuniculus.pro   Lagomorpha
254 ..-...I.......................................D.A...Q.............................D..NVQ.A................L....................... Homo_sapiens.pro
254 ..-...I.......................................D.A...Q.............................D..NVQ.A................L....................... Pan_troglodytes.pro
254 ..-.............................................A.N.Q.............................D..NVQ..................I....................... Macaca_nemestrina.pro
254 ..-.............................................A.N.Q.............................D..NVQ..................I....................... Papio_anubis.pro   Primates
254 ..-.............................................A.N.Q.............................D..NVQ..................I....................... Macaca_fascicularis.pro
254 ..-.............................................A.N.Q.............................D..NVQ..................I....................... Macaca_mulatta.pro
249 NR-.NQ.....................P......H.......E...K.D.N.TK.........M.....P............................................................ Loxodonta africana.pro   Proboscidea
253 .EL.N......................N....YPE.......S..AQKN...MK...T..A..A.I..Y...E...T.......T.N...........M..N.Y.......................E.. Gallus_gallus.pro
253 .EL.N......................N....YPD.......S...QKN...LK...T..A....I..Y...A...T.......T.N...........M..N.Y.......................E.. Meleagris_gallopavo.pro   Galliformes
253 .QF..S.....................P....YPA...........QKN...VK...A..A..S.I..Y...E...K.......T.N...........M..N.Y.......................E.. Anas_platyrhynchos.pro   Anseriformes   Aves
253 PEL..S.....................A....YPA.......S...QKE...VK...A..A..T.I..NA..E...N.......T.N..............N.Y..M....................E.. Aquila chrysaetos chrysaetos.pro
250 PEL..S.....................A....YPA.......S...QKE...VK...A..A..T.I..SA..E...N.......A.N..............N.Y..M....................E.. Haliaeetus albicilla.pro   Accipitriformes
249 .DR..K.....................P....YPD.......NT..EKN.N.SK...A..N....I...P..P...I.......N.-...........M.MK.Y.......................V.. Alligator_sinensis.pro
249 .DR..K.....................P....YPD.......HT..KKN...SK...A..N....I...P..P...N.......N.-...........M.NK.Y.......................V.. Crocodylus_porosus.pro   Crocodilia
253 .DH..S.....................A....YPD.......SE..KKN.N.TK...A..D..M....YK..E...K...I...N.............M..K.Y........S......V.......E.. Pelodiscus sinensis.pro
253 .DR..S.....................A....YPN.......SE..KKN...TK...A..D..I....YK..E...N...I...N.............M..K.Y........S..............E.. Chrysemys picta bellii.pro   Testudines   Reptilia
253 .DR..S.....................A....YPD.......SE..KKN...TK...A..D..I....YE..E...N...I...N.............M..K.Y........S..............E.. Chelonia_mydas.pro
```
