## Supplementary File 1 for "SARS-CoV-2 Spike Glycoprotein and ACE2 interaction reveals modulation of viral entry in wild and domestic animals"

**Note:** Backward stepwise logistic regression model was constructed on all the 32 parameters (29 from FoldX, RMSD, H bonds and delta G) estimated in the 15 known species of infected(11) and uninfected (4). A total of 135 data per parameter (15 hosts x 3 X-ray Crystallography models x 3 selected docking complexes) were analysed.These data were used in seven different combinations based on the combination of X-ray Crystallography structures. The seven combinations include, data against single model - 6LZG,6M0J and 6VW1 (i.e. 45 data); data against two models - 6LZG and 6M0J / 6LZG and 6VW1 / 6M0J and 6VW1 (i.e. 90 data); and data against all the three models - 6LZG and 6M0J and 6VW1 (i.e. 135 data). These seven combinations were evaluated based on the estimates of Number of parameters significant in the logistic model at 1% LS, Number of parameters significant in the logistic model at 5% LS, McFadden's R2, Null deviance, Residual deviance, AIC, p-value of the Chi-sq statistic associated with the null deviance model, p-value of the Chi-sq statistic associated with the residual deviance model, p-value to determine whether there is significant reduction in deviance from null to residual, Hosmer and Lemeshow Goodness of fit (GOF) test (refer to table 2 for details). On evaluating these parameters the best three combinations - 6LZG & 6M0J and 6LZG & 6VW1 and 6M0J & 6VW1& 6LZG, were considered for evaluating the probability in the test data. The test data considered was the known 15 infected and uninfected species. The predicted probability of these 15 species using the selected models is given in Table 2. The predicted probability of all the infected species was closer to being infected with the data combinations - 6M0J & 6LZG followed by 6LZG & 6VW1 and 6M0J & 6VW1& 6LZG. Similar, was the probability for the uninfected species except for a minor difference in *Sus scrofa*. Considering these findings, the combination of 6M0J & 6LZG was selected for predicting the probability of the rest of the species in the study.

**6LZG only**

```
logit <- glm(formula = Infection ~
RMSD+deltaG+Backbone_Hbond+IntraclashesGroup1+Sidechain_Hbond+Solvation_Pol
ar+Solvation_Hydrophobic+electrostatic.kon+energy.Ionisation+Entropy_Complex,
family = "binomial", data = data)
Call:
glm(formula = Infection ~ deltaG + Backbone_Hbond + IntraclashesGroup1 +
    Sidechain_Hbond + Solvation_Hydrophobic + electrostatic.kon,
    family = "binomial", data = data)

Deviance Residuals:
     Min        1Q    Median        3Q       Max
-2.12025  -0.00084   0.00075   0.22321   1.36459

Coefficients:
                      Estimate Std. Error z value Pr(>|z|)
(Intercept)           126.9928    61.0363   2.081   0.0375 *
deltaG                  2.6209     1.3519   1.939   0.0525 .
Backbone_Hbond         -8.2855     4.9095  -1.688   0.0915 .
IntraclashesGroup1     -0.9303     0.5167  -1.801   0.0718 .
Sidechain_Hbond         2.4053     1.6576   1.451   0.1468
Solvation_Hydrophobic   1.6176     0.9051   1.787   0.0739 .
electrostatic.kon     -12.8674     6.6415  -1.937   0.0527 .
---
Signif. codes:  0 '***' 0.001 '**' 0.01 '*' 0.05 '.' 0.1 ' ' 1

(Dispersion parameter for binomial family taken to be 1)

    Null deviance: 52.192  on 44  degrees of freedom
Residual deviance: 15.659  on 38  degrees of freedom
AIC: 29.659

Number of Fisher Scoring iterations: 9

> r2Log <- function(logit) {
+ summaryLog <- summary(backwards)
+ 1 - summaryLog$deviance / summaryLog$null.deviance
+ }
> r2Log(logit)
[1] 0.6999682

> hoslem.test(data$Infection, fitted(logit))

        Hosmer and Lemeshow goodness of fit (GOF) test
```

```
data:  data$Infection, fitted(logit)
X-squared = 0.48303, df = 8, p-value = 0.9999

> 1-pchisq(52.192,44)
[1] 0.1855549
>
> 1-pchisq(15.659,38)
[1] 0.9994944

> 1-pchisq(52.192-21.2703,44-38)
[1] 2.62369e-05
```

**6M0J only**

```
logit <- glm(formula = Infection ~
RMSD+deltaG+IntraclashesGroup1+IntraclashesGroup2+Backbone_Hbond+Sidechain_
Hbond+Van_der_Waals+Electrostatics+Solvation_Polar+Solvation_Hydrophobic+Van_
der_Waals_clashes+torsional_clash+backbone_clash+electrostatic.kon+partial_covale
nt_bonds+energy.Ionisation+Entropy_Complex+Interface_Residues_BB_Clashing,
family = "binomial", data = data)

Call:
glm(formula = Infection ~ RMSD + deltaG + Backbone_Hbond + Van_der_Waals +
    Electrostatics + Solvation_Hydrophobic + Van_der_Waals_clashes +
    electrostatic.kon, family = "binomial", data = data)

Deviance Residuals:
     Min        1Q    Median        3Q       Max
-2.68016  -0.00008   0.11829   0.24732   1.36005

Coefficients:
                      Estimate Std. Error z value Pr(>|z|)
(Intercept)            51.1023    20.1374   2.538   0.0112 *
RMSD                  -52.2261    24.0878  -2.168   0.0301 *
deltaG                  2.5050     1.1514   2.176   0.0296 *
Backbone_Hbond         -3.4859     2.1646  -1.610   0.1073
Van_der_Waals           3.6152     1.8320   1.973   0.0485 *
Electrostatics          1.2791     0.8804   1.453   0.1462
Solvation_Hydrophobic  -2.0457     1.2425  -1.646   0.0997 .
Van_der_Waals_clashes   0.2296     0.1598   1.437   0.1507
electrostatic.kon       4.7013     2.9638   1.586   0.1127
---
Signif. codes:  0 '***' 0.001 '**' 0.01 '*' 0.05 '.' 0.1 ' ' 1

(Dispersion parameter for binomial family taken to be 1)

Null deviance: 52.192  on 44  degrees of freedom
Residual deviance: 19.036  on 36  degrees of freedom
AIC: 37.036

> r2Log <- function(logit) {
+
+  summaryLog <- summary(backwards)
1 - summaryLog$deviance / summaryLog$null.deviance
+ }
> r2Log(logit)
[1] 0.6352702
```

```
> hoslem.test(data$Infection, fitted(logit))

        Hosmer and Lemeshow goodness of fit (GOF) test

data:  data$Infection, fitted(logit)
X-squared = 3.5558, df = 8, p-value = 0.8948
```

**6VW1 only**

```
logit <- glm(formula = Infection ~
RMSD+deltaG+IntraclashesGroup1+IntraclashesGroup2+Backbone_Hbond+Sidechain_
Hbond+Van_der_Waals+Electrostatics+Solvation_Hydrophobic+entropy_mainchain+en
tropy_sidechain+torsional_clash+water_bridge+disulfide+Interface_Residues_VdW_
Clashing+Interface_Residues_BB_Clashing, family = "binomial", data = data)

Call:
glm(formula = Infection ~ RMSD + deltaG + IntraclashesGroup2 +
    Backbone_Hbond + Van_der_Waals + entropy_mainchain + entropy_sidechain +
    torsional_clash + disulfide + Interface_Residues_VdW_Clashing,
    family = "binomial", data = data)

Deviance Residuals:
     Min        1Q    Median        3Q       Max
-1.82292  -0.00002   0.00027   0.08836   1.86144

Coefficients:
                                Estimate Std. Error z value Pr(>|z|)
(Intercept)                    1.016e+03  5.839e+02   1.741   0.0817 .
RMSD                          -5.587e+02  3.311e+02  -1.687   0.0915 .
deltaG                        -2.280e+00  1.439e+00  -1.585   0.1130
IntraclashesGroup2            -2.424e+01  1.386e+01  -1.749   0.0802 .
Backbone_Hbond                 8.155e+00  6.351e+00   1.284   0.1991
Van_der_Waals                  1.428e+00  1.340e+00   1.066   0.2866
entropy_mainchain              1.852e+00  1.464e+00   1.266   0.2056
entropy_sidechain              1.491e+00  1.339e+00   1.113   0.2657
torsional_clash               -1.163e+01  1.057e+01  -1.100   0.2712
disulfide                     -2.372e+15  1.537e+15  -1.543   0.1228
Interface_Residues_VdW_Clashing  1.187e+00  1.090e+00   1.089   0.2762
---
Signif. codes:  0 '***' 0.001 '**' 0.01 '*' 0.05 '.' 0.1 ' ' 1

(Dispersion parameter for binomial family taken to be 1)

    Null deviance: 52.192  on 44  degrees of freedom
Residual deviance: 15.380  on 34  degrees of freedom
AIC: 37.38

Number of Fisher Scoring iterations: 10

> r2Log <- function(logit) {
+ summaryLog <- summary(backwards)
```

```
+
+ 1 - summaryLog$deviance / summaryLog$null.deviance
+ }
> r2Log(logit)
[1] 0.7053305

> hoslem.test(data$Infection, fitted(logit))

        Hosmer and Lemeshow goodness of fit (GOF) test

data:  data$Infection, fitted(logit)
X-squared = 3.4094, df = 8, p-value = 0.9061
```

**6LZG_6MOJ**

```
logit <- glm(formula = Infection ~ RMSD + deltaG + IntraclashesGroup1 +
+ + +     Backbone_Hbond + IntraclashesGroup2 + Sidechain_Hbond + Van_der_Waals
+
+ + +     Electrostatics + Solvation_Polar + Solvation_Hydrophobic +
+ + +     entropy_sidechain + entropy_mainchain + sloop_entropy + torsional_clash +
+ + +     mloop_entropy + backbone_clash + helix_dipole + disulfide +
+ + +     electrostatic.kon + partial_covalent_bonds + Entropy_Complex +
+ + +
Interface_Residues_BB_Clashing+Van_der_Waals_clashes+energy.Ionisation+Numbe
r_of_Residues+Number_of_Residues, family = "binomial", data = data)

Call:
glm(formula = Infection ~ RMSD + deltaG + Backbone_Hbond + IntraclashesGroup2 +
    Sidechain_Hbond + Electrostatics + Solvation_Hydrophobic +
    entropy_sidechain + entropy_mainchain + disulfide + electrostatic.kon +
    Interface_Residues_BB_Clashing + Van_der_Waals_clashes, family = "binomial",
    data = data)

Deviance Residuals:
     Min        1Q    Median        3Q       Max
-2.25951  -0.00467   0.01172   0.29649   2.30808

Coefficients:
                          Estimate Std. Error z value Pr(>|z|)
(Intercept)                1.258e+02  4.019e+01   3.130  0.00175 **
RMSD                      -5.575e+01  2.141e+01  -2.604  0.00325 **
deltaG                     3.636e+00  1.286e+00   2.829  0.00468 **
Backbone_Hbond            -4.571e+00  1.727e+00  -2.647  0.00812 **
IntraclashesGroup2        -1.270e+00  5.215e-01  -2.435  0.01489 *
Sidechain_Hbond            1.821e+00  8.228e-01   2.213  0.02690 *
Electrostatics             1.411e+00  5.894e-01   2.395  0.01663 *
Solvation_Hydrophobic      1.285e+00  4.568e-01   2.812  0.00492 **
entropy_sidechain          8.860e-01  5.018e-01   1.766  0.07746 .
entropy_mainchain         -9.127e-01  5.320e-01  -1.716  0.08625 .
disulfide                 -3.722e+14  1.697e+14  -2.193  0.02830 *
electrostatic.kon         -5.466e+00  2.509e+00  -2.179  0.02936 *
Interface_Residues_BB_Clashing 1.122e+00  6.776e-01   1.656  0.09782 .
Van_der_Waals_clashes      2.513e-01  9.224e-02   2.724  0.00645 **
---
Signif. codes:  0 '***' 0.001 '**' 0.01 '*' 0.05 '.' 0.1 ' ' 1

(Dispersion parameter for binomial family taken to be 1)
```

```
    Null deviance: 104.385  on 89  degrees of freedom
Residual deviance:  36.348  on 76  degrees of freedom
AIC: 64.348

Number of Fisher Scoring iterations: 8
> r2Log <- function(logit){
+ summaryLog <- summary(backwards)
+ 1 - summaryLog$deviance / summaryLog$null.deviance
+ }
> r2Log(logit)
[1] 0.6517876


> hoslem.test(data$Infection, fitted(logit))

        Hosmer and Lemeshow goodness of fit (GOF) test

data:  data$Infection, fitted(logit)
X-squared = 7.6413, df = 8, p-value = 0.00

Infection ~ RMSD + deltaG + Backbone_Hbond + IntraclashesGroup2 +
    Sidechain_Hbond + Electrostatics + Solvation_Hydrophobic +
    entropy_sidechain + entropy_mainchain + disulfide + electrostatic.kon +
    Interface_Residues_BB_Clashing + Van_der_Waals_clashes
```

**6LZG_6VW1**

```
logit <- glm(formula = Infection ~ RMSD + deltaG + IntraclashesGroup1 +
+ + +     Backbone_Hbond + IntraclashesGroup2 + Sidechain_Hbond + Van_der_Waals
+
+ + +     Electrostatics + Solvation_Polar + Solvation_Hydrophobic +
+ + +     entropy_sidechain + entropy_mainchain + sloop_entropy + torsional_clash +
+ + +     mloop_entropy + backbone_clash + helix_dipole + disulfide +
+ + +     electrostatic.kon + partial_covalent_bonds + Entropy_Complex +
+ + +
energy.Ionisation+Number_of_Residues+Interface_Residues+Interface_Residues_Cl
ashing, family = "binomial", data = data)

Call:
glm(formula = Infection ~ RMSD + deltaG + IntraclashesGroup1 +
    Backbone_Hbond + Sidechain_Hbond + Van_der_Waals + Solvation_Polar +
    entropy_sidechain + entropy_mainchain + torsional_clash +
    helix_dipole + disulfide + electrostatic.kon + energy.Ionisation,
    family = "binomial", data = data)

Deviance Residuals:
     Min        1Q    Median        3Q       Max
-2.28819  -0.00001   0.07553   0.36878   2.55569

Coefficients:
                     Estimate Std. Error z value Pr(>|z|)
(Intercept)         4.851e+01  1.432e+01   3.386 0.000709 ***
RMSD               -1.396e+02  5.327e+01  -2.620 0.008788 **
deltaG              1.082e+00  4.532e-01   2.387 0.016989 *
IntraclashesGroup1 -1.948e-01  1.140e-01  -1.708 0.087555 .
Backbone_Hbond     -1.630e+00  7.854e-01  -2.076 0.037920 *
Sidechain_Hbond     7.723e-01  4.280e-01   1.804 0.071165 .
Van_der_Waals       2.000e+00  8.122e-01   2.462 0.013815 *
Solvation_Polar     1.491e+00  4.817e-01   3.096 0.001963 **
entropy_sidechain  -1.217e+00  5.099e-01  -2.387 0.017009 *
entropy_mainchain  -1.093e+00  4.367e-01  -2.504 0.012268 *
torsional_clash    -3.228e+00  1.426e+00  -2.264 0.023559 *
helix_dipole       -2.881e+00  1.675e+00  -1.720 0.085437 .
disulfide          -2.248e+14  1.434e+14  -1.568 0.116809
electrostatic.kon  -3.183e+00  1.582e+00  -2.012 0.044272 *
energy.Ionisation   4.973e+01  3.362e+01   1.479 0.139097
---
Signif. codes:  0 '***' 0.001 '**' 0.01 '*' 0.05 '.' 0.1 ' ' 1

(Dispersion parameter for binomial family taken to be 1)
```

```
    Null deviance: 104.38  on 89  degrees of freedom
Residual deviance:  43.57  on 75  degrees of freedom
AIC: 73.57

Number of Fisher Scoring iterations: 8
>  r2Log <- function(logit){
+ summaryLog <- summary(backwards)
+ 1 - summaryLog$deviance / summaryLog$null.deviance
+
+ }
> r2Log(logit)
[1] 0.5826035

> hoslem.test(data$Infection, fitted(logit))

        Hosmer and Lemeshow goodness of fit (GOF) test

data:  data$Infection, fitted(logit)
X-squared = 3.2232, df = 8, p-value = 0.9196


> formula(backwards)
Infection ~ RMSD + deltaG + IntraclashesGroup1 + Backbone_Hbond +
    Sidechain_Hbond + Van_der_Waals + Solvation_Polar + entropy_sidechain +
    entropy_mainchain + torsional_clash + helix_dipole + disulfide +
    electrostatic.kon + energy.Ionisation
```

**6MOJ_6VW1**

```
> logit <- glm(formula = Infection ~ RMSD + deltaG + IntraclashesGroup1 +
+ Backbone_Hbond + IntraclashesGroup2 + Sidechain_Hbond + Van_der_Waals +
+ Electrostatics + Solvation_Polar + Solvation_Hydrophobic +
+ entropy_sidechain + entropy_mainchain + sloop_entropy + torsional_clash +
+ mloop_entropy + backbone_clash + helix_dipole + disulfide +
+ electrostatic.kon + partial_covalent_bonds + Entropy_Complex +
+ energy.Ionisation+Number_of_Residues+Interface_Residues
+Interface_Residues_Clashing, family = "binomial", data = data)

Call:
glm(formula = Infection ~ RMSD + deltaG + IntraclashesGroup1 +
    Electrostatics + torsional_clash + disulfide + energy.Ionisation +
    Interface_Residues_Clashing, family = "binomial", data = data)

Deviance Residuals:
     Min        1Q    Median        3Q       Max
-2.72877  -0.01066   0.20682   0.47385   1.51821
Coefficients:
                             Estimate Std. Error z value Pr(>|z|)
(Intercept)                 2.478e+01  9.577e+00   2.587  0.00968 **
RMSD                       -5.164e+01  1.807e+01  -2.858  0.00427 **
deltaG                      8.866e-01  4.720e-01   1.879  0.06031 .
IntraclashesGroup1         -1.737e-01  9.106e-02  -1.908  0.05641 .
Electrostatics              1.086e+00  4.047e-01   2.683  0.00730 **
torsional_clash            -2.279e+00  1.190e+00  -1.915  0.05544 .
disulfide                  -3.236e+14  1.242e+14  -2.606  0.00916 **
energy.Ionisation           4.758e+01  2.192e+01   2.171  0.02994 *
Interface_Residues_Clashing 3.291e-01  1.835e-01   1.793  0.07301 .
---
Signif. codes:  0 '***' 0.001 '**' 0.01 '*' 0.05 '.' 0.1 ' ' 1

(Dispersion parameter for binomial family taken to be 1)
    Null deviance: 104.385  on 89  degrees of freedom
Residual deviance:  53.635  on 81  degrees of freedom
AIC: 71.635

Number of Fisher Scoring iterations: 7

>  r2Log <- function(logit){
+ summaryLog <- summary(backwards)
+
+ 1 - summaryLog$deviance / summaryLog$null.deviance
+ }
> r2Log(logit)
```

```
[1] 0.4861802
> hoslem.test(data$Infection, fitted(logit))

        Hosmer and Lemeshow goodness of fit (GOF) test

data:  data$Infection, fitted(logit)
X-squared = 13.521, df = 8, p-value = 0.09513
```

**6LZG_6VW1_6M0J**

```
Call:
glm(formula = Infection ~ RMSD + deltaG + IntraclashesGroup1 +
    Sidechain_Hbond + Electrostatics + Solvation_Polar + Solvation_Hydrophobic +
    entropy_mainchain + backbone_clash + disulfide + energy.Ionisation +
    Number_of_Residues + Interface_Residues_Clashing +
Interface_Residues_VdW_Clashing +
    Interface_Residues_BB_Clashing, family = "binomial", data = data)

Deviance Residuals:
     Min        1Q    Median        3Q       Max
-2.24155  -0.01132   0.13664   0.39175   2.10904

Coefficients:
                              Estimate Std. Error z value Pr(>|z|)
(Intercept)                  3.405e+02  1.859e+02   1.831 0.067049 .
RMSD                        -5.395e+01  1.532e+01  -3.521 0.000430 ***
deltaG                       1.318e+00  4.456e-01   2.957 0.003106 **
IntraclashesGroup1          -1.432e-01  9.206e-02  -1.555 0.119905
Sidechain_Hbond              8.089e-01  3.491e-01   2.317 0.020503 *
Electrostatics               6.031e-01  3.155e-01   1.911 0.055987 .
Solvation_Polar              6.316e-01  1.786e-01   3.536 0.000406 ***
Solvation_Hydrophobic        6.539e-01  2.342e-01   2.793 0.005227 **
entropy_mainchain           -4.097e-01  2.872e-01  -1.426 0.153740
backbone_clash              -7.674e-01  2.599e-01  -2.953 0.003146 **
disulfide                   -2.161e+14  1.113e+14  -1.941 0.052248 .
energy.Ionisation            5.206e+01  2.720e+01   1.914 0.055624 .
Number_of_Residues          -3.874e-01  2.322e-01  -1.668 0.095235 .
Interface_Residues_Clashing      1.906e+00  8.881e-01   2.146 0.031879 *
Interface_Residues_VdW_Clashing -1.817e+00  8.577e-01  -2.119 0.034121 *
Interface_Residues_BB_Clashing   1.841e+00  6.348e-01   2.900 0.003732 **
---
Signif. codes:  0 '***' 0.001 '**' 0.01 '*' 0.05 '.' 0.1 ' ' 1

(Dispersion parameter for binomial family taken to be 1)

    Null deviance: 156.577  on 134  degrees of freedom
Residual deviance:  69.916  on 119  degrees of freedom
AIC: 69.916

Number of Fisher Scoring iterations: 7

> r2Log <- function(logit){
+
```

```
+ summaryLog <- summary(backwards)
+ 1 - summaryLog$deviance / summaryLog$null.deviance
+ }
> r2Log(logit)
[1] 0.5534734

> hoslem.test(data$Infection, fitted(logit))

	Hosmer and Lemeshow goodness of fit (GOF) test

data:  data$Infection, fitted(logit)
X-squared = 5.9379, df = 8, p-value = 0.6542

> formula(backwards)
Infection ~ RMSD + deltaG + IntraclashesGroup1 + Sidechain_Hbond +
    Electrostatics + Solvation_Polar + Solvation_Hydrophobic +
    entropy_mainchain + backbone_clash + disulfide + energy.Ionisation +
    Number_of_Residues + Interface_Residues_Clashing +
Interface_Residues_VdW_Clashing +
    Interface_Residues_BB_Clashing
```
