## Supplementary Table 1 for "SARS-CoV-2 Spike Glycoprotein and ACE2 interaction reveals modulation of viral entry in wild and domestic animals"

**Supplementary Table 1. Species considered in this study**

| **Order** | **Name (common name)** | **Accession number** | **Accession number** |
| --- | --- | --- | --- |
| Artiodactyla | *Bos indicus* **(Indian Cattle)** | XM_019956160.1 | XP_019811719.1 |
|  | *Bos indicus x Bos taurus* **(Indian crossbred Cattle)** | XM_027533926.1 | XP_027389727.1 |
|  | *Bos taurus* **(Exotic Cattle)** | XM_005228428.4 | XP_005228485.1 |
|  | *Bubalus bubalis* **(Buffalo)** | XM_006041540.2 | XP_006041602.1 |
|  | *Bison bison bison* **(American bison)** | XM_010834699.1 | XP_010833001.1 |
|  | *Camelus bactrianus* **(Double humped Camel)** | XM_010968001.1 | XP_010966303.1 |
|  | *Camelus dromedaries* **(Single humped camel)** | XM_010993415.2 | XP_010991717.1 |
|  | *Capra hircus* **(Goat)** | NM_001290107.1 | NP_001277036.1 |
|  | *Ovis aries* **(Sheep)** | XM_012106267.3 | XP_011961657.1 |
|  | *Sus scrofa* **(Pig)** | NM_001123070.1 | NP_001116542.1 |
| Perissodactyla | *Equus asinus* **(Donkey)** | XM_014857647.1 | XP_014713133.1 |
|  | *Equus caballus* **(Horse)** | XM_001490191.5 | XP_001490241.1 |
| Chiroptera | *Pteropus alecto* **(Black fruit bat)** | XM_006911647.1 | XP_006911709.1 |
|  | *Rhinolophus ferrumequinum* **(Greater horseshoe bat)** | AB297479.1 | BAH02663.1 |
|  | *Myotis brandtii* **(Brandt's bat)** | XM_014544294.1 | XP_014399780.1 |
|  | *Eptesicus fuscus* **(Big brown bat)** | XM_008154928.2 | XP_008153150.1 |
|  | *Desmodus rotundus* **(Common vampire bat)** | XM_024569930.1 | XP_024425698.1 |
|  | *Phyllostomus discolor* **(Pale spear-nosed bat)** | XM_028522516.1 | XP_028378317.1 |
|  | *Rousettus aegyptiacus* **(Egyptian fruit bat)** | XM_016118926.1 | XP_015974412.1 |
| Pholidota | *Manis javanica* **(Sunda pangolin)** | XM_017650257.1 | XP_017505746.1 |
| Carnivora | *Felis catus* **(Cat)** | XM_023248796.1 | XP_023104564.1 |
|  | *Panthera tigris altaica* **(Siberian Tiger)** | XM_007090080.2 | XP_007090142.1 |
|  | *Mustela putorius furo* **(Ferret)** | XM_004758885.2 | XP_004758942.1 |
|  | *Canis lupus familiaris* **(Dog)** | NM_001165260.1 | NP_001158732.1 |
|  | *Vulpes vulpes* **(Red Fox)** | XM_025986727.1 | XP_025842512.1 |
|  | *Lontra canadensis* **(North American river otter)** | XM_032880138.1 | XP_032736029.1 |
| Rodentia | *Mus musculus* **(Mouse)** | NM_027286.4 | NP_081562.2 |
|  | *Rattus norvegicus* **(Rat)** | NM_001012006.1 | NP_001012006.1 |
|  | *Cricetulus griseus* **(Hamster)** | XM_027432806.1 | XP_027288607.1 |
| Lagomorpha | *Oryctolagus cuniculus* **(Rabbit)** | XM_002719845.3 | XP_002719891.1 |
|  | *Ochotona princeps* **(American pika)** | XM_004597492.2 | XP_004597549.2 |
| Primates | *Homo sapiens* **(Human)** | NM_001371415.1 | NP_001358344.1 |
|  | *Pan troglodytes* **(Chimpanzee)** | XM_016942979.1 | XP_016798468.1 |
|  | *Papio anubis* **(Baboon)** | XM_021933040.1 | XP_021788732.1 |
|  | *Macaca nemestrina* **(Southern pig-tailed monkey)** | XM_011735203.2 | XP_011733505.1 |
|  | *Macaca mulatta* **(Rhesus monkey)** | NM_001135696.1 | NP_001129168.1 |
|  | *Macaca fascicularis* **(Crab eating monkey)** | XM_005593037.2 | XP_005593094.1 |
| Proboscidea | *Loxodonta Africana* **(African elephant)** | XM_023555192.1 | XP_023410960.1 |
| Galliformes | *Gallus gallus* **(Chicken)** | XM_416822.5 | XP_416822.2 |
|  | *Meleagris gallopavo* **(Turkey)** | XM_019612009.2 | XP_019467554.1 |
| Anseriformes | *Anas platyrhynchos* **(Mallard)** | XM_013094461.3 | XP_012949915.2 |
| Accipitriformes | *Aquila chrysaetos chrysaetos* **(Golden Eagle)** | XM_029999165.1 | XP_029855025.1 |
|  | *Haliaeetus albicilla* **(White-tailed eagle)** | XM_009927339.1 | XP_009925641.1 |
| Crocodilia | *Alligator sinensis* **(Chinese alligator)** | XM_025210843.1 | XP_025066628.1 |
|  | *Crocodylus porosus* **(Salt water alligator)** | XM_019529281.1 | XP_019384826.1 |
| Testudines | *Pelodiscus sinensis* **(Chinese softshell turtle)** | XM_006122829.3 | XP_006122891.1 |
|  | *Chelonia mydas* **(Green sea turtle)** | XM_007070499.1 | XP_007070561.1 |
|  | *Chrysemys picta bellii* **(Painted turtle)** | XM_024108749.1 | XP_023964517.1 |
