## Supplementary Table 2 for "SARS-CoV-2 Spike Glycoprotein and ACE2 interaction reveals modulation of viral entry in wild and domestic animals"

**Supplementary Table 2. Within Mean group distance among the Orders**

| **Order** | **Within Mean group distance (DNA)** | **Within Mean group distance (Protein)** |
| --- | --- | --- |
| Perrisodactyla | 0.01 | 0.02 |
| Primates | 0.02 | 0.03 |
| Accipitriformes | 0.03 | 0.03 |
| Crocodilia | 0.04 | 0.05 |
| Carnivora | 0.07 | 0.10 |
| Testudines | 0.07 | 0.11 |
| Artiodactyla | 0.08 | 0.10 |
| Rodentia | 0.10 | 0.12 |
| Lagomorpha | 0.12 | 0.13 |
| Chiroptera | 0.14 | 0.23 |
| Galliformes | 0.21 | 0.28 |
