## Supplementary Table 3 for "SARS-CoV-2 Spike Glycoprotein and ACE2 interaction reveals modulation of viral entry in wild and domestic animals"

**Supplementary Table 3. Between group distance (between Primates and other groups)**

| **Order** | **Primates (DNA)** | **Primates (Protein)** |
| --- | --- | --- |
| Perrisodactyla | 0.131 | 0.164 |
| Carnivora | 0.162 | 0.199 |
| Pholidota | 0.163 | 0.183 |
| Lagomorpha | 0.165 | 0.197 |
| Rodentia | 0.179 | 0.211 |
| Chiroptera | 0.181 | 0.249 |
| Artiodactyla | 0.186 | 0.241 |
| Proboscidea | 0.189 | 0.237 |
| Testudines | 0.516 | 0.573 |
| Crocodilia | 0.518 | 0.565 |
| Accipitriformes | 0.562 | 0.528 |
| Anseriformes | 0.594 | 0.587 |
| Galliformes | 0.605 | 0.653 |
