## Supplementary Table 8 for "SARS-CoV-2 Spike Glycoprotein and ACE2 interaction reveals modulation of viral entry in wild and domestic animals"

**Parameters that are significant between the infected and the uninfected on Unpaired t-test**

| Parameter | Significance (** - 1% level of significance and * - 5% level of significance) |
| --- | --- |
| RMSD | S** |
| deltaG | S* |
| Van_der_Waals | S* |
| entropy_sidechain | S* |
| Solvation_Hydrophobic | S** |
| IntraclashesGroup1 | S** |
